# An artificial motor protein that walks along a DNA track

**DOI:** 10.1101/2025.03.07.641123

**Authors:** Patrik Nilsson, Neil O. Robertson, Nils Gustafsson, Roberta B. Davies, Chu Wai Liew, Aaron Lyons, Ralf Eichhorn, Cassandra S. Niman, Gerhard A. Blab, Elizabeth H.C. Bromley, Andrew E. Whitten, Anthony P. Duff, Ivan N. Unksov, Jason P. Beech, Peter Jönsson, Till Böcking, Birte Höcker, Derek N. Woolfson, Nancy R. Forde, Heiner Linke, Paul M. G. Curmi

## Abstract

Molecular motors are fundamental to life^1–6^ because of their ability to convert chemical energy into mechanical work, an ability that is conferred by the chemical and structural complexity of their constituent proteins. Scientists have long sought to create artificial protein motors that may reveal insights into how biological motors function. While artificial molecular motors based on small molecules^7^ and DNA^8,9^ have been developed, creating an artificial motor protein has remained an elusive goal in synthetic biology^10^. Here we demonstrate the realization of an artificial protein motor called Tumbleweed (TW) that walks directionally along a DNA track under external control. TW consists of three legs, each with a ligand-gated DNA-binding domain that enables selective interaction with specific sites along a DNA track^11^. Using single-molecule fluorescence assays and a programmable microfluidic device, we show that TW steps directionally along a designed DNA track in response to a defined sequence of ligand inputs. We built our TW molecular walker using a modular approach, combining existing proteins with known properties to achieve emergent motor function, similar to how Nature evolves new proteins. Our design strategy thus offers a a platform for engineering advanced and dynamic protein functionality. Our demonstration of TW walking represents a step toward developing fully autonomous protein motors and opens new avenues for uncovering and leveraging the principles by which biological motors transduce chemical energy into motion.

## Main Text

Molecular motors and machines are essential components of cellular function, performing critical mechanical tasks that enable life. Nature has evolved these macromolecules to transduce chemical energy into mechanical work with remarkable efficiency and precision. The evolution and scope of their sophisticated functions are enabled by the chemical and structural complexity of their constituent proteins. Despite their fundamental importance, understanding how these molecular motors derive their function from structure remains a major challenge in structural biology and biophysics.

What if we could build a functioning artificial motor protein? Doing so would give us precise control over design details and enable us to unravel how motor function emerges from the interplay of its functional parts. Artificial protein motors would also provide mechanistic insight into how biological motors transform energy to work efficiently, opening the way for uncovering new, Nature-inspired, energy-efficient engineering principles that are starkly different to our human-made engines powered by chemical fuel. Designing and realizing such an artificial motor protein is therefore a major goal in synthetic biology and nanotechnology.

Despite significant recent progress, this goal has remained elusive. Major efforts in synthetic molecular systems have yielded small-molecule motors^7^ (which earned the 2016 Nobel Prize in Chemistry) and more sophisticated DNA-based motors and machines^8,9^, but none of the design principles so far have achieved the combination of directionality, autonomy, processivity, speed and efficiency found in natural protein motors. Creating a protein motor requires generating functional modules and integrating them into a coordinated machine. *De novo* protein design has recently created components that could, in principle, be incorporated into motors – such as switchable assemblies^12,13^, two-state hinge proteins^14^ and a coupled axle-rotor protein^15^. However, the outstanding challenge remains to couple and coordinate (flexible) motor components so that surface-binding and directed motion are realized in concert with energy transduction to achieve motor function. Despite several road maps for designing artificial protein motors^11,16–18^, to date, none have been realized.

Here we present the first realization of a functional artificial protein motor. It is created through a modular, bottom-up approach that combines biological and designed protein elements that each lack inherent motor function. Our design strategy overcomes limitations in *de novo* protein design by assembling well-characterized functional modules – including ligand-controlled track-binding feet, connecting legs with appropriate flexibility – to create a clocked-walker protein (which we named Tumbleweed, TW^11^) that moves directionally along a DNA track in a controlled fashion (Fig. 1). Our artificial motor outperforms DNA-based synthetic motors in terms of speed while maintaining comparable processivity. Crucially, our modular approach shows how assembling domains with well-defined properties yields emergent motor function from the synergistic interaction of non-motor components. This strategy can be used to engineer other complex protein systems and it provides insight into how natural protein machines may have evolved. TW represents a step towards the development of a fully autonomous protein motor and introduces a new platform for the design of functional and dynamic nanostructures.

**Fig. 1.**
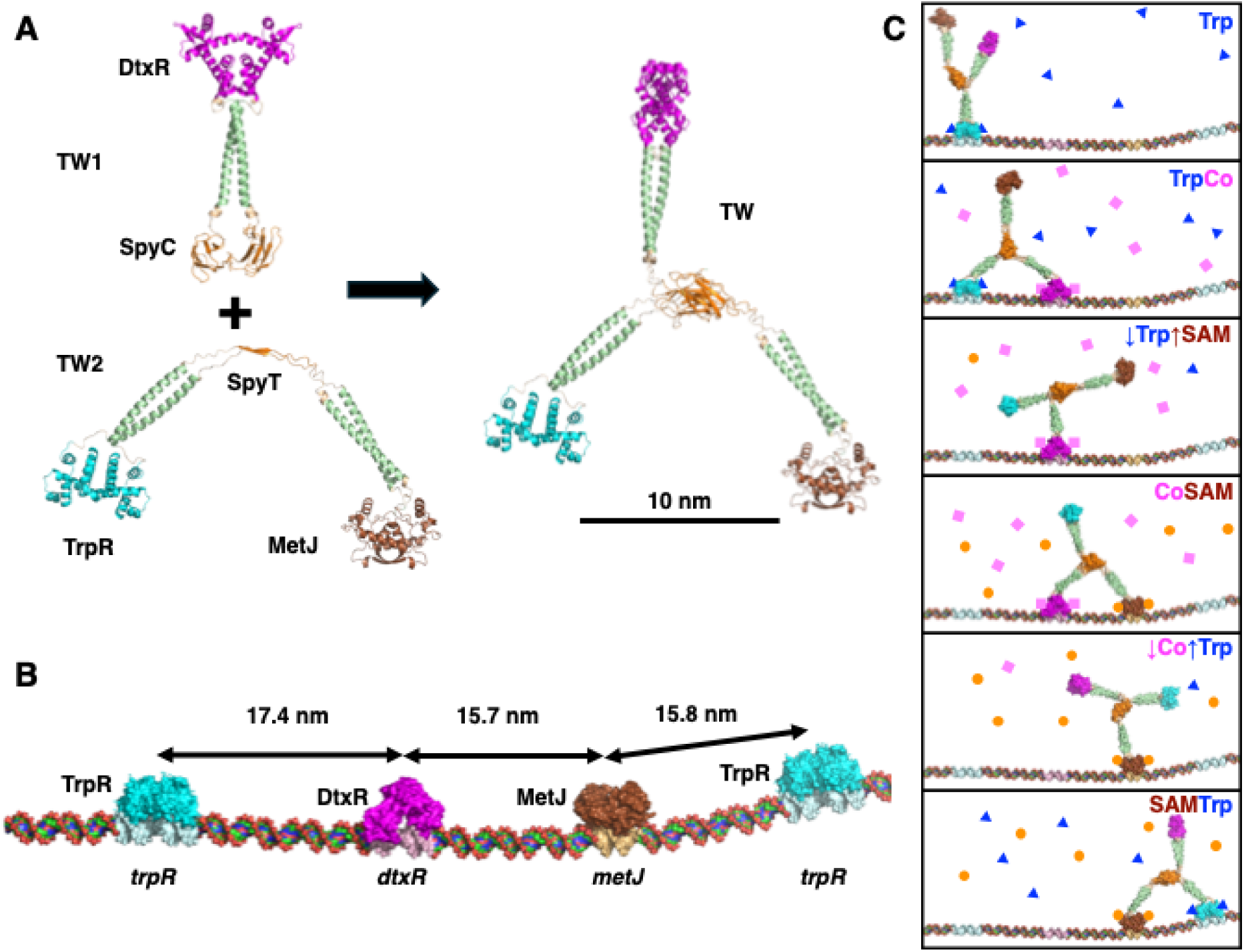
The Tumbleweed (TW) artificial-motor protein concept. (**A**) We constructed TW from two homodimeric proteins, TW1 and TW2. TW1 comprises the DtxR repressor, a geminin helix, and the SpyCatcher protein. TW2 comprises the TrpR repressor, a geminin helix, the SpyTag, another geminin helix, and the MetJ repressor. Combining TW1 and TW2, the SpyTag/SpyCatcher system covalently assembles TW. (**B**) To achieve unidirectional motion, we used a DNA track with an ordered series of cognate binding sites for each of the three TW feet (shown docked): *trpR*-*dtxR*-*metJ*, which can repeat indefinitely, and are arranged so that TW remains on the same face of the track. (**C**) Directional motion was controlled by time-dependent bathing of TW in ligand solutions. In the presence of tryptophan (top panel; Trp, blue triangle), TW is bound by TrpR foot (cyan). Upon adding Co^2+^ (pink squares, second panel from top), the DtxR foot (magenta) also binds to the track adjacent to the TrpR foot. When the concentration of Trp decreases (third panel from top), the TrpR foot releases the track. When adding S-adenosyl methionine (fourth panel from top; SAM; orange circle), the MetJ foot (brown) binds adjacent to the DtxR foot. When Co^2+^ concentration is decreased, the DtxR foot releases (second panel from bottom). Finally, upon re-addition of Trp, the TrpR foot binds adjacent to the MetJ foot (bottom panel). Note that TW and the track were designed so that only one site is reachable for each binding foot (e.g. near but not far *trpR* site in the bottom panel).

### TW modular concept for artificial motor proteins

Our TW concept is a three-legged walker with three key conceptual ingredients^11^ (Fig. 1). First, we designed TW to have three distinct feet, each made from different repressor proteins that bind to cognate DNA sites in the presence of their respective ligand, namely TrpR and its ligand L-tryptophan (Trp), DtxR and Co^2+^, and MetJ and S-adenosyl-L-methionine (SAM). These feet are connected to a central hub by α-helical coiled-coil legs (Fig. 1A).

Second, to ensure unidirectional motion, we designed the DNA track to present specific binding sites for the TW feet in a unique order – *trpR*, *dtxR*, *metJ* (italics with lowercase first letter used to represent cognate sites on DNA), which repeats for the length of the track (Fig. 1B) – which enables TW to walk in this order (from left to right in Fig. 1C). Specifically, we designed the two-leg stride length so that when TW is bound to the DNA track via one foot (for example TrpR bound to *trpR* in the presence of Trp), it can reach and bind to the adjacent site, *dtxR*, when the next ligand, Co^2+^, is introduced (Fig. 1C, top), but not the previous *dtxR* site, which is separated from the TW-bound *trpR* site by an intervening *metJ* site (Fig. 1B).

Third, to control the TW motion we bathed TW in solutions comprising pairs of controlling ligands in a cyclical, time-dependent manner: with (Trp + Co^2+^), then (Co^2+^ + SAM), then (SAM + Trp). By doing so, TW moves from a state where it is, first, bound to the track via TrpR and DtxR, next by DtxR and MetJ, and then MetJ and TrpR (Fig. 1C), before the cycle is repeated. In between these relatively stable, two-foot-bound states, TW will transit more vulnerable, single-foot-bound states (Fig. 1C), where there is a higher probability of detaching from the DNA track. The direction of motion can be controlled by altering the order in which ligand solutions are presented.

The externally controlled, time-dependent bathing solution thus supplies TW with the free energy to power its unidirectional motion by controlling ligands and their chemical potentials. Each DNA-binding module will bind its respective ligand when the concentration of the ligand is high (high chemical potential) and will eventually dissociate and release its controlling ligand when the ligand concentration is low (low chemical potential). Because each of the TW’s steps is enabled by diffusive motion, TW is a Brownian ratchet^19^.

### Realizing the TW concept to achieve directed motion

#### Constructing the three-legged TW

Because three-legged, heterotrimeric proteins are uncommon in nature, we designed and assembled our three-legged TW using the SpyTag/SpyCatcher system^20^, in two parts: TW1 and TW2. We produced each by expressing synthetic genes in *Escherichia coli* and purifying the His-tagged proteins using immobilized metal affinity chromatography (IMAC; Figs 1A and 2A). The resulting TW1 comprised DtxR, the α-helical domain from the geminin coiled-coil, and SpyCatcher, each connected by flexible Gly-Ser linkers. In turn, TW2 comprised TrpR, the geminin coiled-coil α helix, SpyTag, a second copy of the geminin coiled-coil α helix and MetJ, each connected by Gly-Ser linkers.

**Fig. 2.**
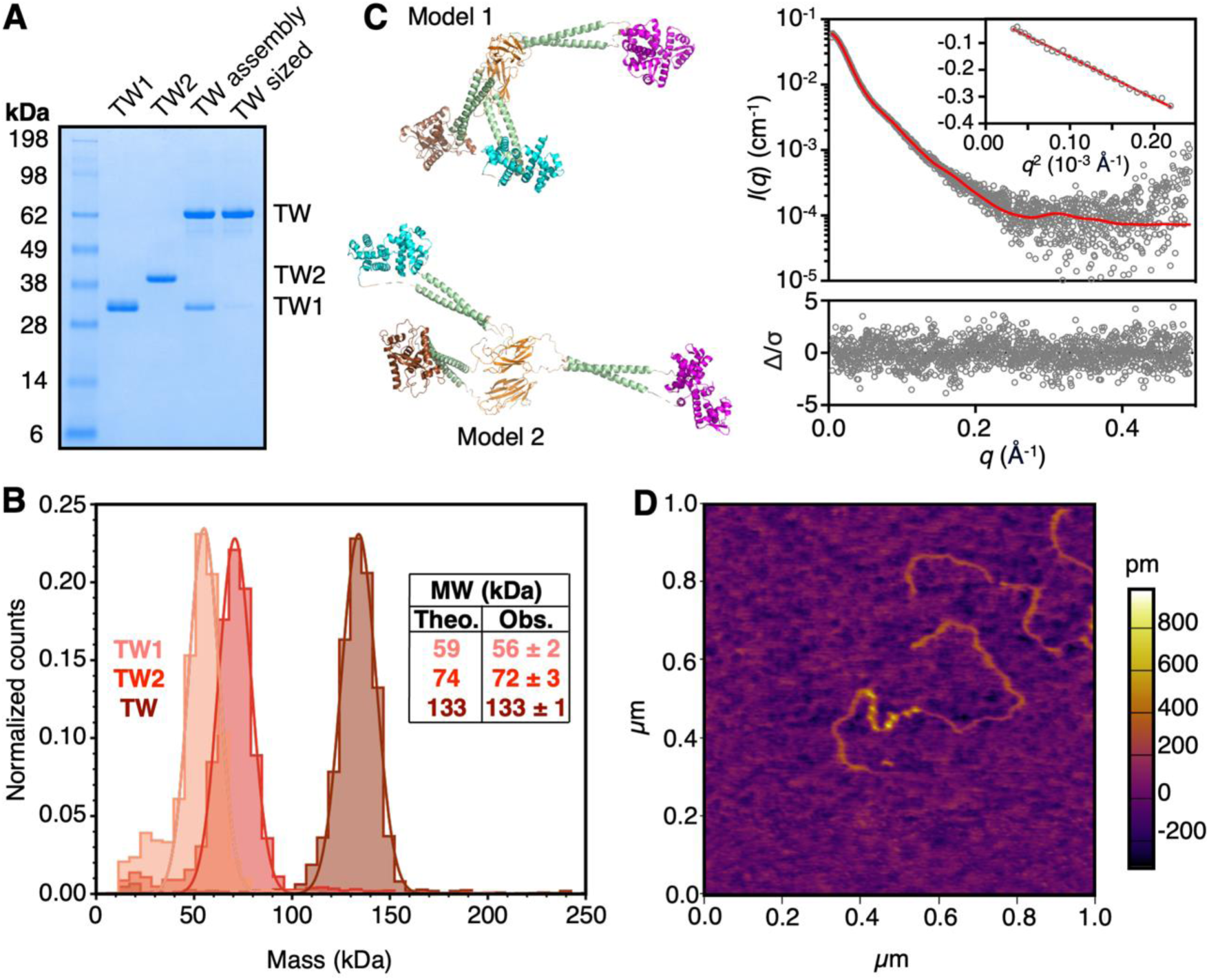
Assembly and characterization of TW. (**A**) SDS-PAGE showing the covalent TW assembly and purified TW after size exclusion chromatography (TW sized). TW sized protein was used for all remaining experiments. (**B**) Mass photometry characterization of purified TW1, TW2 and TW. Insets are the theoretical and observed molecular weights (mean ± SEM, *n* = 4) for each protein species. (**C**) SEC-SAXS analysis of TW. Top right shows the SAXS intensity as a function of *q* (momentum transfer in Å^-1^). Inset shows a Guinier plot of the data; data were fit to an ensemble of models using the program Multi-FoXS. The best fit (top right, red line) as determined by χ^2^ was obtained using two conformers, one L-shaped (Model 1, top left) and one extended (Model 2, bottom left), weighted at 0.725 and 0.275, respectively. The error-weighted residual plot (lower right panel) shows no significant systematic deviations between the model and the experimental data. (**D**) AFM image showing eight TWs bound in the presence of Trp to eight repeats of a *metJ*-*trpR*-*dtxR* track inserted between flanking sequences.

TW1 and TW2 formed homodimeric proteins that were purified to homogeneity (Fig. 2A). We used mass photometry to determine the size of each molecule, which we found to be consistent with the expected mass (Fig. 2B; Extended Data Table 1). Mature TW was then generated by mixing TW1 and TW2, with a slight excess of TW1 (1.2:1). We confirmed the covalent TW product by SDS-PAGE (Fig. 2A). The product was purified by size-exclusion chromatography (SEC) and the size determined by mass photometry, which again was found to be consistent with the expected mass (Fig. 2B; Extended Data Table 1).

#### Characterizing the TW structure

To verify the structural integrity of TW, we collected SEC-coupled small-angle X-ray scattering (SEC-SAXS) data using the Australian Synchrotron (Fig. 2C). The observed radius of gyration (*R_g_* Guinier 67.7 ± 0.6 Å; *R_g_ P*(*r*) 69.1 ± 0.4 Å) was consistent with an extended protein (Extended Data Fig. 1). Given the engineered hinge flexibility of TW, we anticipated an ensemble of structures in solution. We fit the experimental data using Multi-FoXS^21^, where we allowed the program to bend AlphaFold2-generated models^22^ of TW via six hinge points linking the three legs to both the SpyTag/SpyCatcher hub and to each of the DNA binding modules. We obtained a best fit with two conformers: an L-shaped conformer (72.5%; Model 1, Fig. 2C top left) and an extended conformer (27.5%; Model 2, Fig. 2C bottom left). Adding further conformations only marginally improved the fit of the model to the data and the new conformers clustered with the original two.

#### Designing the DNA tracks

We designed DNA tracks to space the binding sites, *trpR*, *dtxR* and *metJ*, such that each protein foot would bind on the same side of the DNA (Fig. 1B). We set the spacing between adjacent sites so that two feet of TW could bind adjacent sites but the stride of TW was not long enough to bind simultaneously to non-adjacent sites (assuming a straight DNA track). We designed the spacing between *trpR* and *dtxR* sites slightly longer because both TrpR and DtxR binding sites are centered on the minor groove of DNA, whereas MetJ binding is centered on the major groove (Fig. 1B).

#### Visualizing TW on an eight-repeat track

Based on the above design principles, we produced a track with eight copies of *metJ*-*trpR*-*dtxR* inserted into the pYIC plasmid^23^. We linearized the resulting track, mixed it with TW in the presence of 10 mM Trp, and visualized it using atomic force microscopy (AFM) after drying on a mica substrate. Images revealed eight bright, evenly spaced features along the DNA (Fig. 2D). The contour lengths of the DNA flanking the eight bright features (representing bound TWs) are consistent with the location of the inserted (*metJ*-*trpR*-*dtxR*)_8_ track.

#### Optimizing binding specificity of TW to DNA

For TW to function as intended, we needed to ensure that each of its feet only bind to its cognate DNA sites with high affinity (*K*_d_ 1-100 nM) and only with its respective controlling ligand. To achieve this, we optimized both the intervening sequences in the DNA track and the solution conditions (controlling ligand concentrations, pH and ionic strength) for binding by characterizing the binding of TW to various DNA tracks (Fig. 3A; Figs. S2-S4) using surface plasmon resonance (SPR).

**Fig. 3.**
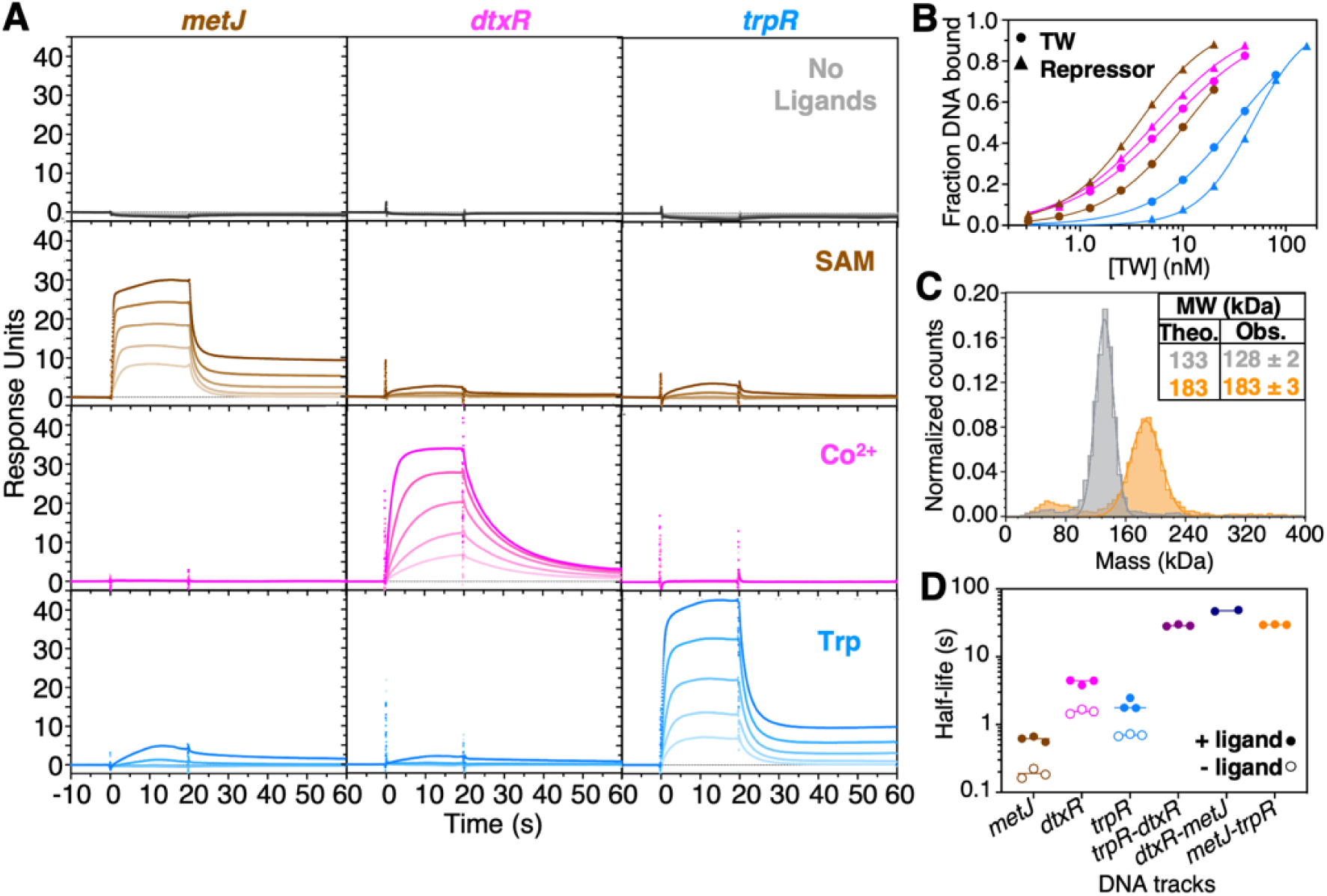
Optimizing TW-DNA track interaction. (**A**) Panel of SPR sensorgrams titrating 5, 10, 20, 40 and 80 nM TW onto DNA oligonucleotides containing single cognate sites (*metJ*, *dtxR* and *trpR* sites) in the presence/absence of controlling ligands show the intended TW binding of each foot (in the presence of ligands) to the respective cognate DNA, with minimal offsite non-specific binding. (**B**) Representative binding curves fitting the Hill equation to average steady state SPR responses. Curves show binding to single site tracks containing individual cognate sites (brown – *metJ*; magenta – *dtxR*; cyan – *trpR*) for both TW (circles) and individual repressor dimers (triangles) in the presence of the controlling ligand. **(C)** Mass photometry data comparing the distribution of estimated molecular masses for the complex of TW and a duplex DNA containing the *metJ* and *trpR* sites (TW:*metJ-trpR* complex) in the absence of ligands (gray) and in the presence of SAM and Trp (orange). The estimated molecular mass of 128 ± 2 kDa in the absence of ligands, and 183 ± 3 kDa in the presence of SAM and Trp are in excellent agreement with the predicted masses of 133 and 183 kDa, respectively. (D) Apparent half-lives of different TW:DNA complexes as calculated from fitted *k*_off_ values determined by SPR experiments in which TW was bound to DNA in the presence (filled circles) or after removal (open circles) of ligands. Dissociation was triggered by adding an excess of free competing specific DNA in the presence or absence of ligands. Controlling ligands correspond to the respective DNA tracks, as indicated. Circles represent the individual values from replicate experiments.

SPR sensorgrams for TW binding to DNA oligonucleotides containing single cognate binding sites showed that, in the absence of ligands, TW bound minimally to DNA (Fig. 3A, top row). However, when we added each ligand, we found the TW to bind to the appropriate cognate DNA site as intended, with minimal offsite non-specific binding (Fig. 3A bottom three rows). From the sensorgrams we calculated the dissociation constants, *K_d_*, for TW to single-site DNA oligonucleotides, which were found to be in the 10–30 nM range (Fig. 3B; Extended Data Table 2). TW dissociation constants were very similar albeit greater than those of purified repressor proteins DtxR and MetJ, while TW had a slightly stronger affinity for *trpR* oligonucleotides in the presence of Trp compared to purified TrpR repressor (Fig. 3B, Extended Data Fig. 2, Extended Data Table 2).

We determined the specificity of each TW foot for its cognate DNA site by titrating TW in the presence of a single controlling ligand on an SPR chip with two-site DNA tracks (Extended Data Fig. 3). In the absence of ligands, TW did not bind to any two-site track where TW was varied from 2.5 to 40 nM (Extended Data Fig. 3A). In this concentration range, no non-specific binding was observed to a track containing *metJ* and *trpR* when Co^2+^ was present (Extended Data Fig. 3C). A low level of non-specific binding was observed in the presence of SAM on a track containing *trpR* and *dtxR* sites when [TW] > 20 nM (Extended Data Fig. 3B). The highest degree of non-specific binding by TW was observed in the presence of the Trp ligand (Extended Data Fig. 3D). Titration of TW onto a track containing *metJ* and *dtxR* sites showed clear, specific binding of TW in the presence of appropriate ligands, either SAM or Co^2+^ (Extended Data Fig. 3B,C,E), while increasing levels of non-specific binding were observed in the presence of non-cognate ligand Trp (Extended Data Fig. 3D,E). However, the apparent affinity of off-target binding to the *dtxR-metJ* track in the presence of Trp was more than threefold lower than the on-target affinities (Extended Data Table 2), and for concentrations of TW below 10 nM, non-specific binding to this track was minimal compared with specific binding (Extended Data Fig. 3D,E).

We confirmed that we obtained the intended stoichiometry of TW:DNA complexes using mass photometry. TW bound all single cognate binding sites in a 1:1 fashion (Extended Data Fig. 5A, Extended Data Table 1). Importantly, TW also formed a 1:1 complex with DNA containing two cognate sites in the presence of both corresponding ligands, even when the DNA was added in 4–10 fold molar excess to TW (Fig. 3C, Extended Data Fig. 5B, Extended Data Table 1). In these experiments, we used 25 nM TW with up to 250 nM DNA, indicating TW preferentially binds on a single DNA molecule at least up to the nanomolar concentration range.

#### Identifying the timescales for optimal binding kinetics

To achieve the envisioned directional motion of the TW clocked walker by sequential binding and release of the feet^11,16,24,25^, it is crucial to ensure appropriate timescales for these processes. For our TW concept, the time that TW spends in a particular two-ligand solution condition must be short compared to the half-life of the corresponding two-foot-bound state, while the release of single feet when ligands are removed must be fast compared to this timescale.

To determine these timescales, we studied the dissociation kinetics of TW:DNA complexes using an SPR displacement assay, where during the dissociation phase, a large excess of free specific DNA was introduced to prevent rebinding of TW to surface-bound DNA. This enabled us to determine apparent off-rates in different ligand conditions (Fig. 3D, Extended Data Fig. 4, Extended Data Table 3). We found the half-lives for TW following the removal of controlling ligands to be < 2 s. In the presence of single controlling ligands, the half-lives were ∼ 0.6 – 4 s. In both the absence and presence of controlling ligands, the MetJ foot was the fastest to dissociate from DNA, and the DtxR foot the slowest. The off-rates decreased dramatically in the presence of two controlling ligands, with half-lives in the ∼30–50 s range. These measurements provided us with the boundaries for clocking our TW to achieve the desired motor function.

### Demonstrating TW stepping along a DNA track

Having assembled the TW and achieving the designed features and functions of each of its constituent components, we next examined its ability to walk along the DNA track using single-molecule Förster resonance energy transfer (FRET). We demonstrate that TW can take multiple steps forward and backward along a DNA track in response to microfluidically controlled ligand exchanges. Modelling based on these results predicts unidirectional motion along an infinite track.

#### Using single-molecule FRET to observe TW stepping

To probe the walking motion, we attached a short DNA track (Fig. 4A) containing four binding sites (from bottom to top: *trpR*, *dtxR, metJ,* and *trpR*) to a PLL-g-PEG passivated surface using a biotin-streptavidin linker. We labelled the track’s two *trpR* sites with two different FRET acceptors, ATTO647N (red emission) at the top and ATTO565 (yellow emission) at the bottom. The corresponding FRET donor, AlexaFluor 488 (green emission), was used to label the TrpR foot of TW close to the DNA binding site (Ser107) to enable FRET when the TrpR foot binds to either of the two *trpR* sites. This enabled us to uniquely identify each of the three different, two-foot-bound, states by the wavelength of their emission. Starting closest to the coverslip we expected: yellow FRET emission with TW bound to the bottom-*trpR* site and the *dtxR* site (Fig. 4B); green emission (*i.e.*, no FRET) with TW bound to the *dtxR* and *metJ* sites (Fig. 4C); and red FRET emission with TW bound to the *metJ* and top-*trpR* sites (Fig. 4D).

**Fig. 4.**
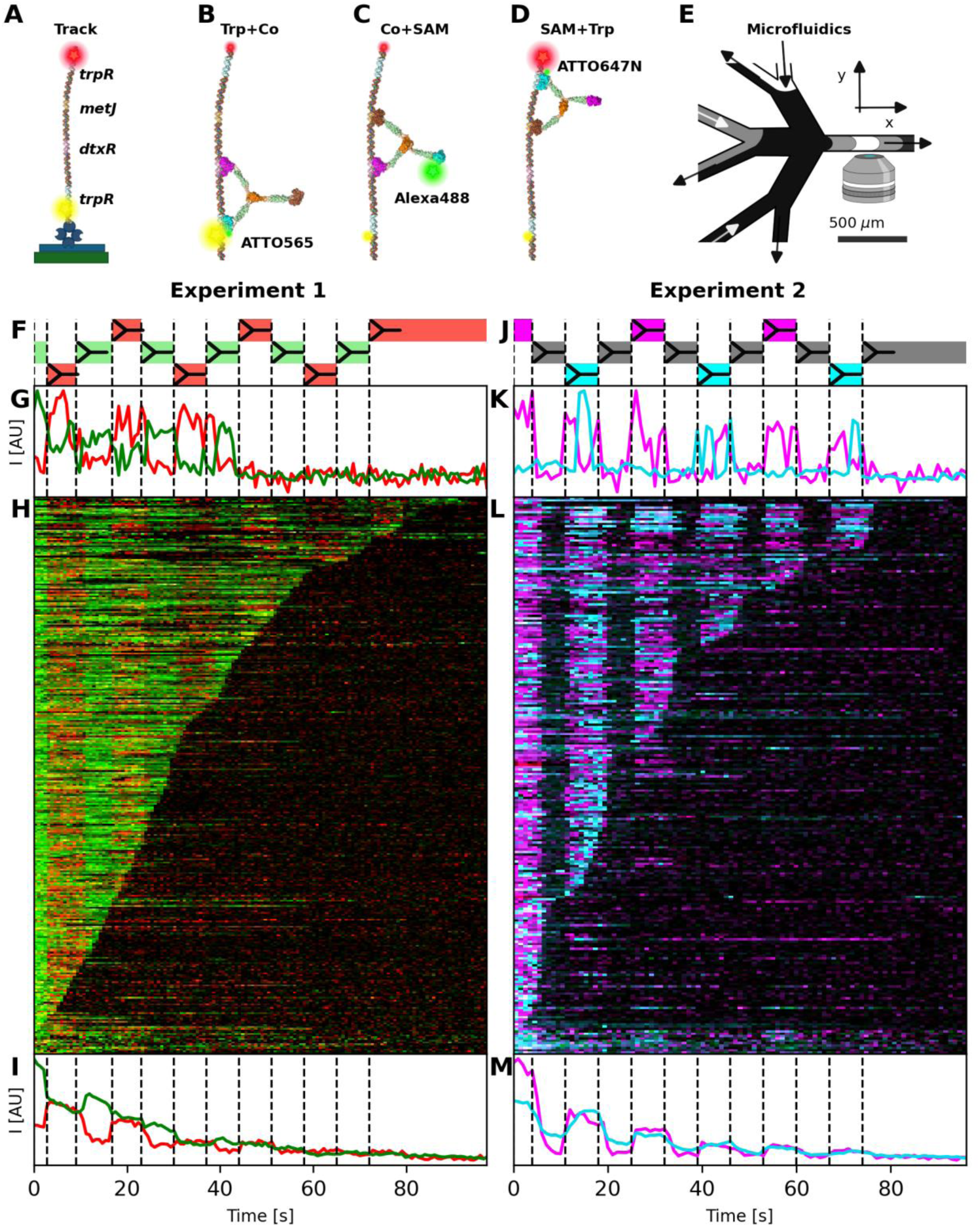
Two experimental demonstrations of TW stepping on a DNA track using single molecule FRET. **(A)** TW walked a four-site (*trpR-dtxR-metJ-trpR* bottom to top) DNA track attached via biotin-streptavidin linker to a biotinylated PLL-g-PEG passivated surface. The *trpR*-sites at the bottom and top were labelled with different FRET acceptors (yellow emitter Atto565 and red emitter Atto647N, respectively). The TrpR foot was labelled with the FRET donor AlexaFluor488, a green emitter. **(B-D)** TW:DNA complexes that we expected to form in presence of pairs of ligands. (Trp + Co^2+^ in **(B)**, SAM + Co^2+^ in **(C)**, SAM + Trp in **(D)**). Each of the three complexes was designed to yield a distinct fluorescence signal on excitation with a 488 nm laser: (**B**) FRET to Atto565 (yellow emission) when the TrpR foot is bound to the bottom-*trpR* site**, (C)** AlexaFluor488 (green emission) from the free TrpR foot, and **(D)** FRET to Atto647N (red emission) when TrpR is bound to the top-*trpR* site. **(E)** Microfluidic device used to rapidly change solution conditions in the imaging volume. **(F-I) Experiment 1**: two-color imaging with emission split at 543 nm to simultaneously record AlexaFlour488 emission (green) and ATTO565 plus ATTO647N emission (red). **(F)** Schematic of expected TW position and false color representation of the fluorescence signal expected when changing solution (dashed lines). **(G)** Normalized example of single-molecule trace. **(H)** Kymographs of all detected colocalizing single molecules sorted by time of last identified fluorescence. **(I)** Sum of all single-molecule traces in the respective experiment. **(J-M) Experiment 2**: same as in (**F-I)**, but using two-color imaging with emission split at 633 nm to simultaneously record ATTO565 emission (FRET emission from the bottom *trpR* site, shown in cyan) and ATTO647N emission (FRET emission from the top *trpR* site, shown in magenta).

To control TW stepping by changing the ligand concentrations in real time, we used a custom-designed microfluidic device (Fig. 4E). This enabled us to switch between three different controlling solutions with minimal drag in arbitrary temporal order, with a time resolution of less than ∼ 0.2 s (ref ^26^).

Ligand-pair solutions were changed at 7 s intervals, which was long enough to collect several fluorescence frames during each ligand step and to allow the slowest foot, DtxR, to dissociate (Extended Data Fig. 4; Extended Data Table 3). Once one foot detached from the track, the TW diffused within a larger volume around the remaining bound foot. This enabled the next foot in the step series to get close enough to bind to the next cognate binding site on a subsecond timescale^24,25^. Given that two-foot-bound states have a half-life of ∼30 s (Extended Data Fig. 4; Extended Data Table 3), we expected to observe an average of > 4 steps per TW molecule.

#### Demonstrating TW stepping: Experiment 1

We report two types of experiment (Fig. 4F-I and Fig. 4J-M, respectively) that together enabled us to fully characterize TW stepping up and down the four-site track. To generate stepping, we sequentially changed the ligand-pair solutions to induce TW to step down the track, then two steps up, then two steps down, and repeating for a total of 11 steps as indicated in Figs 4F and 4J. In each experiment, DNA:TW complexes were identified by colocalization, and time traces of background-subtracted fluorescence intensity from hundreds of individual complexes in a field of view were extracted.

Experiment 1 shows that the TrpR foot of TW binds and unbinds the *trpR* sites as the ligands are changed by the fluidic device, while TW remains attached to the DNA track (Fig. 4F-I), thereby demonstrating that the TW walks along the track. We added 1 nM TW to the fluidic device in the presence of Co^2+^+SAM ligands to induce TW to bind to the central *dtxR* and *metJ* sites of the DNA (Fig. 4C). We collected light simultaneously in two wavelength bands, the first suitable to observe green emission from the TW TrpR foot and the second suitable for emission from both red and yellow fluorophores. This enabled us to observe both FRET positions (at the bottom and the top of the track) in the same channel and detect the signal from the TW as either donor emission or FRET from either acceptor. We observed fluorescence changes in synchrony with solution changes, and how the donor (shown in green in Fig. 4G, H)) and acceptor (shown in red in Fig. 4G, H) emission signals alternated out of phase, as shown in the example trace (Fig. 4G) until the signal was lost through bleaching or full detachment of TW. Overlaid kymographs for each of the two observation channels (Fig. 4H) from one single field of view show that there are 322 complexes where the FRET state of the TW:DNA complex is alternating in synchrony with the change in controlling solution, indicating stepping. The sum of these individual traces (Fig. 4I) further demonstrates shows the same anti-correlated behavior in phase with the changes in controlling solution (with the reduced total signal over time again being attributed to bleaching and TW dissociation). Taken together, Experiment 1 confirms that the TrpR foot binds and unbinds to/from the *trpR* sites on the track as dictated by the ligands. However, it cannot discriminate between TrpR binding to the intended *trpR* site versus the *trpR* site at the opposite end of the track, hence, we conducted Experiment 2.

#### Demonstrating TW stepping: Experiment 2

In Experiment 2 (Fig. 4J-M) we confirmed that TW successfully visits both the top and bottom of the track, as intended, when externally controlling the ligands. To achieve this, we simultaneously imaged two-color emission of the bottom and top FRET acceptors recorded in separate channels. Examples of the anti-correlated FRET signals from *trpR* sites on the top and bottom of the track are shown in Fig. 4K and Extended Data Fig. 6 (magenta = top of track; cyan = bottom of track, see Fig. 4J). From overlaid kymographs (Fig. 4L) for each of the two observation channels from four separate fields of view, we found 236 complexes that exhibited dynamic behavior correlated with the controlling ligand exchange. The dark bands in these kymographs coincide with periods where TW is expected to be bound to the central *metJ* and *dtxR* sites (Fig. 4C). The sum of the individual traces (Fig. 4M) shows that TW starts at the top *trpR* site, moves down through the dark central site and continues to step. The alternating dark and bright states confirm stepping, and the presence of magenta and cyan signals confirm that TW reaches both the top and the bottom of the tracks.

From individual traces (Fig. 4K; Extended Data Fig. 6), we found that when the controlling solution contains Trp, single TWs have a highly dynamic behavior, switching on a timescale of seconds between being bound to the top-*trpR* and the bottom-*trpR* sites (Fig. 4K, L; Extended Data Fig. 6). This indicates that in these experiments TW can ‘overstep’, by binding to the non-adjacent *trpR* site. By analyzing all traces during the first three solution conditions (Fig. 4L; Extended Data Fig. 6), we found that the TW TrpR foot bound the intended (adjacent) *trpR* site ∼65% ± 7.7% of the time (correct stepping), whereas it bound the unintended, distal *trpR* site ∼35% ± 6.3% of the time (overstepping). Importantly, as we show below in our modeling, the motor moves forward directionally, on average, regardless of such overstepping.

#### Control experiments

We also performed control experiments that show that no stepping by TW on the four-site oligonucleotide track occurs when switching solutions with the same ligands (keeping ligand concentrations constant while microfluidically switching between identical solutions), confirming that stepping is caused by changing ligand concentrations, and that any hydrodynamic forces due to the microfluidics play no role in the observed motion (Supplementary Discussion A, Extended Data Fig. 7). When ligands are removed, we observe that TW falls off the tracks as expected (Supplementary Discussion A, Extended Data Figs 8-9).

#### Coarse-grain modeling predicts directional motion along infinite track

To understand how the observed stepping behavior translates into TW’s ability to walk along a long DNA track that contains many *trpR - dtxR - metJ* repeats, we created a coarse-grained model based on Master equations governing the binding and unbinding events between the three TW feet and the track (Supplementary Discussion B). Based on the experimentally observed binding and unbinding rates (Extended Data Table 3), and including the observed likelihood to overstep, we predict that a TW on an infinite track would move directionally at a speed of about 0.5 track periods per full solution cycle (Extended Data Fig. 10), or ∼1.2 nm/s using the 21 s experimental cycle time. The processivity is ∼ 50 – 100 nm (Extended Data Figs 11 and 12) and is governed by the observed detachment time in the presence of two ligands.

## Discussion

We created the first artificial motor protein, a long-standing goal in synthetic biology and protein design. By employing a modular, bottom-up approach combining well-characterized protein domains, we demonstrated that motor function can emerge from the strategic assembly of non-motor components. We showed how our three-legged TW artificial protein motor – a clocked Brownian walker that binds specifically to its track as a monomer with nanomolar affinity – moves directionally and controllably on a DNA track when time-dependent external supply of ligands controls how each foot alternatingly binds to its cognate DNA site.

The processivity of TW observed in the single molecule FRET experiments is consistent with the two-foot bound half-life, measured in the SPR experiments. From the kymographs (Figs 4H and L), we deduced a bound half-life of ∼30-40 s (Fig. 4H and L), which agrees well with the expected bound time of a TW based on the dissociation rates measured in SPR. The kymographs are sorted by time of the last observed fluorescence signal; this occurs when the TW either bleached or dissociated from the track. Because our experiments were designed to make maximum use of the available photon budget during the expected bound time, the two possible events (bleaching and dissociation) occur on similar timescales in our experiments and are thus indistinguishable. The coincidence of the observed TW track-bound half-life with that of the three possible two-foot-bound states (Extended Data Table 3) indicates that TW only transiently populates the more vulnerable single-foot-bound states during ligand exchange.

Our single-molecule experiments show that individual TWs successfully take up to 11 steps up and down a short track before dissociating. The dynamic behavior also indicates that in the presence of the Trp ligand, TW can ‘overstep’, sometimes binding to the non-adjacent *trpR* site instead of the intended *trpR* site that is adjacent to the bound foot. Most likely this overstepping occurs because of inherent flexibility of the DNA track (Supplementary Text 3) and the TW linker regions. Similar dynamic, stochastic behavior, including occasional misstepping, is well known from single-molecule studies of biological motors ^27–29^. The main consequence of overstepping is variability in the displacement of individual TWs (Fig. S11) which is insensitive to the precise probability of overstepping over a wide parameter range (Figs S10-S12). In the absence of overstepping, which may be achievable by using DNA tracks based on rigid origami nanotubes^30^, our modelling predicts that TW approaches a speed of one-track period per solution cycle. This corresponds to 2.3 nm/s using the 21 s experimental cycle time and may be sped up by reducing the solution cycle time. Our motor outperforms artificial DNA motor speed (typically 10^-2^ - 1 nm/s) with comparable processivity. While speed is also limited by the timescale for detachment of TW feet, a speed of 10 – 20 nm/s is likely achievable, bringing it closer to the performance of natural protein motors.

By creating a protein walker from non-motor components, we have shown that a modular protein engineering approach can deliver a new, designed function. This accomplishment reduces the size of synthetic protein-based motors from modular assemblies of tens to thousands of proteins^31,32^ to individual motor proteins constructed from modular domains. Our approach circumvents fundamental limitations in current protein design capabilities – namely, the difficulty in creating proteins with the required dynamic properties for motor function – whilst mimicking how new functional proteins evolve in Nature by combining and assembling existing protein domains or protein fragments to render emergent properties^33^. Our proof-of-principle demonstration is evidence that complex, dynamic protein machines can be developed without requiring complete mastery of designing flexible, information-transmitting protein structures from scratch.

Our clocked walker serves as an experimental platform for investigating the physical principles underlying nanoscale motor function, including the trade-offs between energy efficiency and performance predicted by stochastic thermodynamics. It also establishes a critical foundation for developing fully autonomous motor proteins. We propose a roadmap organized around three critical elements that must be integrated to achieve autonomy: the enzymatic cycle to harness free energy, allosteric communication to couple binding and enzymatic states, and structural asymmetry to ensure directionality^16^. Each of these properties can be found individually in natural proteins, and the engineering of each represents a significant but achievable challenge. Natural non-motor proteins that already exhibit some of these properties could serve as starting points for engineering. Alternatively, existing proteins could be modified to introduce these capabilities, or completely new proteins could be designed *de novo* with the specific functions in mind. The ultimate challenge lies in integrating all three properties into a single, coherent design – a goal that, while ambitious, now appears within reach given our successful demonstration of a modular motor protein.

Taken together, our creation of the first artificial motor protein founds a strategy for the design and fabrication of functional molecular devices that leverages the atom-scale accuracy of protein design as well as the highly scalable and sustainable methods of bioproduction. As the field advances toward autonomous motor proteins and their integration into larger systems, these molecular machines could be applied, for example, in energy-efficient, network-based computing, in active materials, and other forms of sustainable nanoengineering.

## Methods

### Construct design and cloning

Protein constructs were based on the following proteins: DtxR (UniProt Acc. No. P0DJL7), Geminin (UniProt Acc. No. E2QRF9), TrpR (UniProt Acc. No. P0A881), MetJ (UniProt Acc. No. P0A8U6) and the SpyTag:SpyCatcher system^34^. For the repressor and Tumbleweed (TW) constructs DtxR refers to residues 2—121 with a C102D mutation (based on PDB: 1F5T^35^), Geminin_Coiled-Coil_ refers to residues 110—145 with an additional N-terminal threonine and C-terminal glutamine (based on PDB: 1T6F^36^), TrpR refers to residues 2—108, and MetJ refers to residues 2—105 with a Q45K mutation (based on PDB: 1MJM^37,38^).

Both MetJ and TrpR were expressed from pET28a vectors as fusion proteins with a N-terminal 8 x His-tag. DtxR was expressed from a pET19b vector as a fusion protein with a C-TW was produced from two separate constructs, TW1 and TW2 (Extended Data Fig. 13). TW1 consists of DtxR—Geminin_Coiled-Coil_—SpyCatcher. TW2 consists of TrpR— Geminin_Coiled-Coil_—SpyTag—Geminin_CoiledCoil_—MetJ. For both TW1 and TW2 the protein domains were connected using flexible glycine-serine-rich linkers. Both TW1 and TW2 constructs were synthesized by Genescript within a pET15b vector allowing for the expression of protein with an N-terminal 8 x His-tag and TEV protease cleavage site.

### Protein expression and purification

All proteins were expressed in *E. coli*. Individual repressors and TW1 were expressed using BL21 cells in 1–2 L of LB media. TW2 was poorly expressed under these conditions. After screening expression in different cell and media types TW2 was eventually expressed using C41 cells in 6–12 L of 2xTY media. Cells were grown with agitation at 37 °C until the culture reached OD_600_ ∼ 0.6, at which point the incubation temperature was reduced to 24 °C and protein expression was induced by the addition of IPTG to a final concentration of 0.4 mM. Cultures were then incubated with agitation for approximately 20 h, after which cells were harvested by centrifugation and pellets stored at –80 °C until further processing.

Cell pellets were resuspended at 4 °C with stirring in 20 mM Tris pH 8, 150 mM NaCl, 20 mM Imidazole, 10% glycerol, 2.5 mM MgCl_2_, 0.5 mM CaCl_2_, 1 mM NaN_3_, 0.1 mg/mL lysozyme, 10 µg/mL DNase I and 1 x protease tablet (Sigma-Aldrich). Approximately 25 mL of lysis buffer per liter of cell culture was used to resuspend the cell pellets. The resuspended cell slurry was then lysed using a cell disruptor at 20 kPSI pressure. The soluble fraction was harvested by centrifugation by 50 000 x g and then loaded onto a 5 mL HisTrap^TM^ HP nickel affinity chromatography column (Cytiva). The column was then subjected to a series of washes using a combination of buffer A (20 mM Tris pH 8, 400 mM NaCl, 10% glycerol, 1 mM NaN_3_) and buffer B (buffer A + 1 M imidazole); first with 2% buffer B, then 5% buffer B, and then finally an elution step with 30% buffer B.

TW1 and TW2 were subjected to TEV cleavage and reverse His-Trap purification to increase the purity of the final product. After the initial His-trap purification, the eluted proteins were concentrated and diluted to a final concentration of 3% buffer B. This sample was then subjected to an overnight TEV protease cleavage (1:50 mg/mg TEV:protein), with DTT added to the sample to a final concentration of 1 mM. The TEV-cleaved sample was then flowed over the HisTrap^TM^ column to remove contaminants, collecting the flowthrough.

Finally, all proteins were subjected to size-exclusion chromatography (SEC) using either a HiLoad Superdex 75 26/60 column for individual repressors, or a HiLoad 200 16/600 column for TW1 and TW2 (Cytiva). SEC was performed in 20 mM Tris pH 8, 400 mM NaCl, 10% glycerol, 1 mM NaN_3_ and 1 mM EDTA. Protein yields per cell culture volume were typically ∼ 1 mg/L for individual repressors, ∼4 mg/L for TW1 and ∼0.3 mg/L for TW2. Purified protein was then frozen using liquid nitrogen and stored at –80 °C until further processing.

### Assembly and purification of Tumbleweed

Purified TW1 and TW2 were mixed with a molar ratio of 1.2:1 (with TW2 at a final concentration of ∼20–40 µM) in 20 mM Tris pH 8, 400 mM NaCl, 10% glycerol, 1 mM NaN_3_ and 1 mM EDTA at 4 °C for 16 h. The assembly mixture was then subjected to SEC using a Superdex 200 Increase 10/300 column (Cytiva), using the same buffer as the assembly. Purified assembled Tumbleweed was then frozen in liquid nitrogen and stored at – 80 °C until further processing.

### Fluorescent labelling of TW

Fluorescent TW was produced by thiol conjugation of a maleimide dye to a TW containing a cysteine mutation. A TW2 construct containing TrpR_S107C_ (henceforth referred to as TW2_S107C_) was synthesized by Genscript. The protein was expressed and purified as described above, with 1 mM DTT included in purification buffers to maintain the protein in a reduced state.

Alexa Fluor™ 488 C_5_ Maleimide (AF488; Thermofisher Scientific) was resuspended to 10 mM in DMSO. Purified TW2_S107C_ was reduced with 1 mM TCEP pH 7 prior to labelling at a final concentration of 30–60 µM with 5–10x molar excess of maleimide dye. Reactions were performed in 50 mM Tris pH 8, 400 mM NaCl, 10% (v/v) glycerol either for 2 h at room temperature or overnight at 4°C. Reactions were quenched with 50 mM DTT for 30 min at room temperature. TW1 was added to the mixture such that the final molar ratio of TW1:TW2 was 1.2:1, and left to assemble TW as described above. The assembly mixture was purified from excess dye by SEC using a Superdex 200 Increase 10/300 column (Cytiva), in the same buffer as the assembly.

Following removal of excess dye, the degree of protein labelling was measured using the UV-Vis function of an ND-1000 Nanodrop Spectrophotometer (ThermoFisher Scientific). The absorbance of the protein:dye conjugate was measured at 280 nm (protein) and at 495 nm (AF488) (See Extended Data Table 4). The degree of labelling was calculated according to the equation^39^:

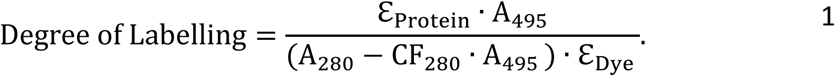

Here, ℇ_Protein_ is the extinction coefficient of the protein at 280 nm (55350 M^-1^ cm^-1^), ℇ_Dye_ is the extinction coefficient of AF488 at 495 nm (73000 M^-1^ cm^-1^) and CF_280_ is the correction factor for the absorption of light at 280 nm by AF488 (0.11). TW was assessed to be ∼100% labelled (two labels per dimeric TrpR foot), and frozen with liquid nitrogen and stored at - 80°C.

### SAXS data collection

Small-angle X-ray scattering (SAXS) data were collected on the small/wide angle X-ray scattering beamline at the Australian Synchrotron^40^, part of ANSTO in Melbourne, Australia. The camera length used was 2680 mm corresponding to a *q*-range of 0.005 – 0.50 Å^-1^.

Proteins were auto-loaded from a 96-well plate and 50 mL of ∼10 mg/mL of sample injected onto a Superdex 200 Increase 5/150 GL equilibrated in 10 mM HEPES pH 7.4, 150 mM NaCl, 5mM MgCl_2_, connected in line with a coflow cell^41^ through which X-rays were passed at 8 x 10^12^ photons per second at 11.5 keV. The data was collected on a Pilatus 2M detector (Extended Data Table 5).

### SAXS data processing

SAXS data were processed using software from the ATSAS suite^42,43^ (Extended Data Table 5). Protein elution profiles were generated from scattering intensities using custom written scripts. The elution profiles were used to determine the signals corresponding to protein and buffer frames. Firstly, 21 buffer frames were selected and averaged for buffer subtraction.

The radius of gyration (*R_g_*) was then determined for each frame across the protein peak (with buffer subtraction performed for each individual frame). Guinier analysis to determine *R_g_* was performed in PRIMUS^42^ to monitor data quality. Distance distribution (*P(r*)) curves were generated using GNOM^42^ to estimate the maximum dimension (*D*_max_) of the protein.

### Multi-state modelling with MultiFoXS

Multi-state modelling using MultiFoXS^21^ was used to determine the population-weighted conformational states of TW that could be contributing to the experimental SAXS profile. The inputs used for MultiFoXS included a protein model, SAXS profile, flexible hinge residues and rigid body connections. To obtain protein models to use as templates for MultiFoXS, models of TW1 and TW2 were generated using ColabFold v1.4: AlphaFold2 using MMseqs2^44^. These Alphafold-generated models were subsequently used to construct a complete model for TW assembly using UCSF Chimera^45^. For MultiFoXS, TW was divided into rigid elements and hinge regions. The rigid elements comprise: the three repressor dimers, the three coiled-coils and the SpyTag/SpyCatcher hub. Six hinges were placed in the Gly-Ser rich linkers. To simplify the computation by MultiFoXS, the polypeptide chain was cut in one of the two Gly-Ser rich linkers joining each rigid element. Thus, the six hinges were defined as: TW1: Gly125-Gly126 and Gly170-Gly171, between DtxR, the coiled-coil and SpyCatcher, respectively; and TW2: Ser114-Gly115, Ser155-Ser156, Gly183-Ser184, Gly228-Gly229, between TrpR, the coiled-coil, SpyTag, coiled-coil and MetJ, respectively (see Extended Data Fig. 13).

Preliminary tests were initially conducted with conformational sampling of 100 conformations and final calculations were performed to sample 10,000 conformations. For each sampled conformation, a SAXS profile is calculated followed by scoring of multi-state models to generate population-weighted ensembles with associated *R_g_*. The best multi-state models that fit the experimental SAXS data were based on the *χ^2^* values and residual plots.

### Design and assembly of DNA tracks

The *metJ* cognate sequence (GAGACGTCTC) was taken directly from the consensus Met box sequence from the MetJ:DNA crystal structure (PDB: 1MJM^37^). The *trpR* cognate sequence (GTACTCGCTAGCGAGTAC) was based on a *trpR^S^* sequence previously shown to bind TrpR dimer with a 1:1 stoichiometry^46^. The *dtxR* cognate sequence (TTAGGTTAACCTAA) was based on a DtxR:DNA crystal structure (PDB: 1F5T^35^), with the sequence altered to produce a palindrome that is capable of binding only one DtxR dimer.

Flanking sequences were designed to create blocks that could be rearranged within a multisite track that maintains consistent sequences either side of the cognate sites. The *metJ* flanking sequences were designed based on the native *E. coli metC* protomer sequence (GenBank Ref. Seq. NC_000913.3.). This promoter sequence was chosen on the basis that it contains a single Met box^37^. The *dtxR* flanking sequences were conserved from the DtxR:DNA crystal structure (PDB: 1F5T^35^). The *trpR* flanking sequences were designed by a genetic algorithm to minimize the presence of cryptic repressor binding sites^23^. Double, triple and quadruple site tracks were designed by arranging the cognate sites and their flanking sequences. Models of the tracks were produced using cgNA+web^47,48^ and analyzed using PyMOL^49^ to ensure the cognate sites were appropriately distanced from one another and remained in the same phase (i.e. on the same side of the track).

All tracks were ordered as single strand oligonucleotides from Integrated DNA Technologies (IDT). Oligonucleotides were resuspended to 100 µM in Milli-Q water and stored at –20 °C. Complementary oligonucleotides were annealed at concentrations of up to 20 µM in 25 mM HEPES pH 7.4, 200 mM KCl, 5 mM MgCl_2_ by first heating the sample to 95 °C for 2 min, and then subjecting to a 95–20 °C gradient at 1°C/min. Annealed tracks were stored at -20 °C until further processing. Extended Data Table 6 lists all DNA tracks used in SPR and mass photometry experiments.

### Atomic force microscopy (AFM)

*E. coli* containing the pK8 plasmid^23^ were cultured in LB media overnight in the presence of 25 mg/mL kanamycin. They were pelleted by centrifugation for 3 min at 12000 rpm. The plasmid was purified using a MiniPrep kit (Qiagen). The DNA was linearized by digestion with EagI (nine parts purified plasmid to one part NEBuffer 3, with 1 unit of EagI restriction endonuclease for every 150 ng of DNA) for 2 h at 25°C, after which the enzyme was heat inactivated by incubation at 65 °C for 25 min. The product was purified using a PCR Cleanup kit (Qiagen). Completion of restriction digest was confirmed by gel electrophoresis.

TW (0.26 mg/ml) and 10X HEPES buffer (40 mM HEPES, 100 mM NaCl, 20 mM MgCl_2_, pH 7.4) were both diluted five-fold in Milli-Q water (1:1:3 TW:10X HEPES:H_2_O). 2.5 µL of this stock was placed into a pre-lubricated microcentrifuge tube, to which was added 1.25 µL of linearized DNA (69.3 ng/mL) and 1.25 µL of 40 mM tryptophan. This mixture was incubated at room temperature on a spinner for 1 hour. After this time, 45 µL of 1X HEPES Buffer was added to the incubated solution, mixed with a pipette, and plated immediately onto freshly cleaved mica, where it was allowed to sit for 4 minutes. Following this, the excess solution was shaken off, and the plate was rinsed five times with 1 mL of Milli-Q water, then dried gently under a stream of filtered compressed air to remove any residual liquid.

AFM images were recorded using an Asylum Research MFP-3D SPM, using Mikromasch HQ:NSC15/AL BS probes with a spring constant of 40 N/m and resonance frequency of 325 kHz.

The chain-tracing software SmarTrace^50^ was used to determine the contour lengths of DNA flanking the bound TW proteins. For this analysis, *N*=5 images were chosen that included seven or eight TW bound to clearly define the inserted track cassette. The cassette is located asymmetrically with respect to the ends of the linearized plasmid, and thus the long and short contour lengths, and their ratio of lengths, were used to confirm that the TW were bound in the expected region of the DNA.

### Surface Plasmon Resonance (SPR)

SPR experiments were performed on an Biacore^TM^ S200 instrument (Cytiva). All experiments were performed in running buffer (25 mM HEPES, pH 7.4, 200 mM KCl, 5 mM MgCl_2_, 0.2 mM EDTA, 0.05% (v/v) Tween-20), which was passed through a 0.22 µm filter and degassed by vacuum for at least 30 min prior to use. Data was collected at 40 Hz in multi detection mode, with the instrument temperature set to 25 °C. All sensorgrams were double referenced.

DNA coating of SPR chips was prepared as previously described^51^. Series S CM5 sensor chips (Cytiva) were initially prepared by depositing ∼5000 RU of streptavidin on to the chip surface via amine coupling. A biotinylated single-stranded oligonucleotide (henceforth referred to as the biotin anchor) was added to the chip for a response of ∼50–70 RU. Finally, annealed single or double cognate site DNA containing a single-stranded overhang complementary to the biotin anchor was flowed over the non-reference channels for a response of ∼10 RU. Chips were regenerated using injections of 10 mM glycine pH 2.5, which removed the annealed DNA and made the biotin-anchor re-available to bind new DNA.

Individual repressors and TW were titrated onto single and double site DNA at 100 µL/min. When required, ligands were used with the following concentrations: 1 mM SAM, 0.4 mM CoCl_2_ and 0.5 mM Trp. All titrations were performed using the A-B-A injection method, with a 30 s initial ligand injection, a contact injection containing protein for 15–30 s, and a final ligand injection of 40–60 s to trigger dissociation. A 60 s injection of buffer containing 0.5 M KCl was used to remove any residual protein from the surface before the next titration point. The average steady state positions were determined by the Biacore Evaluation software (Cytiva) and fit to a Hill equation using Prism 10 (Graphpad):

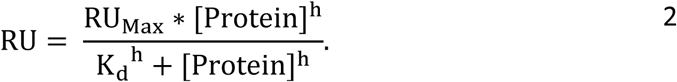

Here, *K*_d_ is the apparent affinity, h is the Hill coefficient and RU_Max_ is the fitted maximum binding response.

We were unable to fit the above titrations to a simple 1:1 kinetic model. Increasing the amount of DNA on the chip surface lead to sensorgrams with slower apparent binding kinetics, indicating the system suffered from mass transfer effects^52^. Protein:DNA interactions are particularly prone to mass transfer in SPR as electrostatically steered interactions lead to high association rates^53^. A consequence of mass transfer is that during the dissociation phase of an SPR experiment, the protein can release and rebind to the surface before it enters the bulk solution, resulting in a slower apparent dissociation rate^53,54^.

To avoid complications arising from TW rebinding DNA on the SPR surface, we decided to study TW:DNA dissociation kinetics using a SPR-based displacement assay which was based on classic tracer experiments^55,56^. This involved dissociating TW in the presence of free DNA in solution, such that TW would preferentially bind to the free DNA and not rebind to the surface. A dual injection method was used to first load TW onto the surface DNA with an immediate second injection used to trigger dissociation. The assay was performed using a 100 µL/min flow rate. For single-site DNA experiments, a 5 nM TW solution was used to bind TW to DNA on the SPR chip in the relevant ligand for 25 s, before dissociation of the TW:DNA complex was triggered with an immediate 60s injection of a solution containing 1 µM free DNA ± ligand. The free DNA contained the relevant cognate binding site but lacked the single stranded anchor overhang and was thus unable to bind the sensor chip. For experiments involving double-site DNA, 1 nM TW was loaded onto the DNA in the presence of two ligands for 60 s before dissociation was triggered with a 60 s injection of a solution containing both ligands and 1 μM of each of the relevant free DNA sequences. The resulting dissociation curves were fit to a single exponential decay function using Prism (Graphpad):

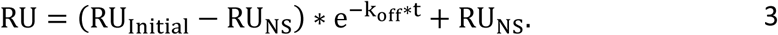

Here, RU_Initial_ is the initial response value, RU_NS_ is the non-specific response value, and k_off_ is the apparent dissociation rate constant. For double-site DNA, double-ligand experiments, RU_NS_ was set to 0.

### Mass photometry

Mass photometry experiments were performed as described^57^. Experiments were performed on an Refeyn TwoMP instrument using 24 x 50 mm high precision 1.5H coverslips (Marienfeld) and CultureWell^TM^ gaskets (Grace Biolabs). Coverslips were first prepared by sonicating in a 50% (w/w) isopropanol bath for 3 min, followed by another 3 min sonication in MilliQ-Water. These sonication rounds were repeated for a total of 3 rounds before the coverslips were dried with nitrogen gas and stored at room temperature. Gaskets were affixed to the coverslip immediately prior to an experiment.

Buffers (25 mM HEPES, pH 7.4, 200 mM KCl, 5 mM MgCl_2_, and 0.2 mM EDTA ± 0.5 mM SAM, 0.4 mM CoCl_2_ and/or 0.5 mM Trp) were passed through a 0.22 µm filter prior to use in mass photometer experiments. Data were collected over a 60 s timeframe using the normal measurement mode and regular image size as set by the Refeyn AcquireMP software. Mass calibrations were performed using a combination of purified bovine serum albumin (Sigma-Aldrich) and NativeMark^TM^ unstained protein standard ladder (ThermoFisher Scientific), with the 66, 132 and 480 kDa peaks providing a calibration curve with R^2^ > 0.99. Manually picked mass distributions within the relevant mass ranges were selected and fit with a Gaussian function to determine an average molecular mass using the Refeyn DiscoverMP software. SAM buffer was found to produce an interference with the mass photometry signal. Additionally, DNA transiently interacted with the glass surface when in the presence of CoCl_2_. Both conditions result in an additional mass peak centered around 60–80 kDa.

TW1, TW2 and TW samples were added to slides to a final concentration of 10 nM. TW and 4–10-fold molar excess DNA were preincubated for ∼5 min before analysis on the mass photometer. Final concentrations of 25–30 nM TW and 250–300 nM DNA were used for single-site DNA experiments, whereas double-site DNA experiments used 5–25 nM TW and 50–250 nM DNA.

### DNA tracks for single molecule observation

Single strand oligonucleotides used to generate DNA tracks (Extended Data Table 7) for single molecule experiments were ordered from IDT. Oligonucleotides were resuspended to 100 µM in Milli-Q water and stored at –20 °C. All oligonucleotides required for the track were annealed at concentrations of 11.4 µM of each component oligo in 70 µL 1.25X PNK (T4 polynucleotide kinase reaction) buffer (New England BioLabs, Cat. No. B0201S) by first heating the sample to 95 °C for 2 min, followed by a 95—20 °C gradient at 1°C/min. The annealed oligonucleotides were then ligated by adding 8 µL ATP (New England BioLabs, Cat. No. P0756S) to a final concentration of 1 mM and 2 µL T4 DNA ligase (New England BioLabs, Cat. No. M0202S) to a final concentration of 10,000 units/ml. The resulting 80 µL ligation reaction mixture in 1X PNK was kept at 16 °C for 16 h before heat inactivation at 65°C for 10 min. The DNA was purified using bind-wash-elute to a commercial spin column (QIAquick PCR Purification Kit, Qiagen) as per manufacturer’s instructions. Purified tracks were stored at 4 °C until use. Models of the tracks were produced using cgNA+web^47,48^ and analyzed using PyMOL^49^ to ensure the cognate sites were appropriately distanced from one another and remained in the same phase.

### Microfluidics for single molecule observation

To switch the TW running solutions, a previously published microfluidic device, developed by Niman *et al.*^26^, was used (Fig. 4E; Extended Data Fig. 14). This device is designed to allow switching between three solutions (three combinations of ligands) in arbitrary time order, with sub-second time resolution, and without stopping the flow. It consists of three inlet channels of equal length that intersect at the entrance to a primary observation channel where the TW stepping takes place. Each inlet has an “escape” channel of equal length, microfluidic resistance and pressure originating directly before the intersection and terminating at a common waste outlet (circle, Extended Data Fig. 14A far left). These escape channels provide routes away from the observation channel. Increasing the flow rate in any one of the three inlet channels propagates that solution alone along the observation channel, while pushing the other two solutions away from the observation channel to their respective escape channels. This design allows for the solution in the observation channel to be changed rather than stopped, allowing for faster and more complete switching, as well as for switching the three fluids in arbitrary time order. Flow rates were controlled using Fluigent lineup pumps, flow unit and OxyGEN software (Fluigent). Flow rates of 7 µL/min in the “on” channel and 1 µL/min in the “off” channels provided good switching and arbitrary sequences of ligand solutions were achieved by the application of 7 µl/min to any of the inlet channels (and 1 µl/min to the others) (Extended Data Fig. 14C).

The time resolution for switching the ligand concentration in immediate proximity to the channel surface where TW is located (the chemical switching time) is limited by the diffusion time of ligands in and out of the non-slip boundary layer. The chemical switching time inside the boundary layer depends on the fluid speed and the location along the microfluidic channel^26^. For the experiments reported here, observations were conducted in the main observation channel, approximately 1 mm away from the intersection with the inlet lines. A 3D finite element simulation (incompressible, laminar flow) of the volumetric flow rates in the device was performed using COMSOL Multiphysics software with the fluid set as water, with a density of 1000kg/m^3^ and a dynamic viscosity of 0.001 Pa s. For inlet flow rates of 7, 1 and 1 µl/min, the simulated volumetric flow rate in the observation channel of 2.285 µl/min which corresponds to a fluid speed in the observation channel of 7.6 mm/s. Based on the results in ^26^ (Extended Data Fig. 14) we expect that the chemical switching time in the experiments reported here was about 0.2 s or less, much faster than the 7 s during which each solution was held constant.

### Microfluidic master mold fabrication protocol

Microfluidic devices were produced by fabricating a mold in which polydimethylsiloxane (PDMS) could be cured and subsequently bonded to a glass surface. To fabricate the mold, a photomask was fabricated using an MLA150 maskless lithography system (Heidelberg Instruments GmbH). A 4” silicon wafer was dried at 180°C for 5 min. A 50 µm thick dry film resist (SUEX®, K50, DJ Microlaminates) was applied to the wafer using a laminating machine (Catena 35, Acco UK Ltd) at 65 °C. A pre-exposure bake was performed at 85 °C on a hotplate (Model 1000-1 Precision Hot Plate, Electronic Micro Systems Ltd) to remove air bubbles and to relax the film. SUEX film was exposed at 365 nm in a contact mask aligner (Karl Suss MJB4 soft UV) for 27 s at a lamp power of 30 mW/cm^2^. Post exposure baking was done at 85 °C for 5 min. Development was done in Mr DEV 600 (Micro Resist Technology GmbH) for 15 min plus 5 min in fresh developer followed by rinsing in flowing isopropyl alcohol and drying with nitrogen. A final bake was done in a convection oven at 200 °C for 15 min. To reduce adhesion of PDMS to the master, a layer of aluminum oxide (∼1 nm) followed by a monolayer of perfluordecyltriklorosilan (FDTS) was deposited in an atomic layer deposition system (Fiji – Plasma Enhanced ALD, Veeco).

### Microfluidic device preparation and single molecule observation

Base imaging buffer (25 mM HEPES, pH7, 200 mM NaCl, 5 mM MgCl_2_) was passed through 0.22 µm filter and stored at 4 °C for no longer than 1 month.

Coverslips (40 mm x 20 mm, #1.5 thickness, VWR) were prepared by sonicating in isopropyl alcohol (Merck) for 10 min and blow dried with pressurized nitrogen. These were then cleaned with acid piranha solution of 3 parts sulfuric acid (Merck) and 1 part hydrogen peroxide (Merck) for 30 min at 80 °C after which they were rinsed with MilliQ water (Millipore MQ) and blow dried with pressurized nitrogen. Final cleaning was performed in oxygen-nitrogen mixed plasma (Zepto Plasma Cleaner, Diener Electronic) for 3 min at 40 kHz, 100W.

PDMS microfluidic devices were prepared by thoroughly mixing 2-component Sylgard 184 silicone elastomer and curing agent (G A Lindberg) in a 10:1 ratio and degassing to remove air bubbles. The mixture was poured onto the device master mold and allowed to cure at 80°C for > 1h. After curing, using a 1 mm biopsy tool, inlets were punched at the three points indicated by inward pointing arrows (Extended Data Fig. 14A). Using a 3 mm biopsy tool, outlets were punched at the circles (Extended Data Fig. 14A) indicating the common waste outlet and the 10 mm mark along the primary observation channel. The molded PDMS was washed with isopropyl alcohol. Immediately after coverslip cleaning, the molded PDMS and coverslip contact surfaces were both etched in nitrogen plasma for 10 s and then bonded together by placing the molded PDMS onto the cleaned coverslip. Immediately, the device was wetted and then incubated for a minimum of 1h in imaging buffer with 1 mg/ml of 1:10000 PLL-g-PEG-Biotin (3.4 kDa PEG):PLL-g-PEG (2 kDa PEG). The device was visually inspected via white light transmission microscopy at 20X magnification for blockages, bonding defects, and bubbles. Devices were stored at RT submerged in imaging buffer.

Before use, the devices were washed with imaging buffer by aspirating from the common waste outlet while filling the observation channel outlet with buffer under 1 atmosphere pressure. The device channels were then incubated in imaging buffer containing 20 nM streptavidin (Merck) for at least 10 min. Glass slides were cut to size and glued to the ends of the coverslip using two component dental cement (Abberior Instruments) (Extended Data Fig. 14B). Immediately before imaging, 0.2 mg/ml TROLOX (Merck) was added to the imaging buffer which was bubbled with nitrogen for 15 min to remove dissolved oxygen.

The microfluidic system was prepared by washing all microfluidic lines and flow meters with chlorine, MilliQ water, isopropyl alcohol (Merck) and finally with MilliQ water. The cleaned microfluidic system was loaded with three 2 ml reservoirs containing the ligand-pair solutions; imaging buffer, 2 mg/mL TROLOX (Merck) and

1. 0.5 mM Trp, 0.2 mM CoCl_2_
2. 0.2 mM CoCl_2_, 1mM SAM
3. 1mM SAM, 0.5 mM L-Trp.

For each control experiment, the microfluidic system was loaded with two reservoirs containing the same ligand-pair solution, either 1, 2, or 3, and the third reservoir containing imaging buffer, 2 mg/mL TROLOX (Merck) and no ligands.

Finally, the device was connected to the microfluidic system via the three 1 mm inlets and rinsed with running flows set to 7 µl/min for the solution desired in the observation channel and 1 µl/min for the two alternate solutions. After rinsing for a minimum of 1 min the device was mounted and secured to the microscope.

Single molecule fluorescence imaging was performed using a Nikon-TI2 inverted optical microscope in TIRF mode. A 100 × TIRF objective (Plan-APOCHROMAT 100 × 1.45 NA Oil, Nikon) was used to collect fluorescence onto an sCMOS camera (Prime 95B, Photometrics), yielding a pixel size of 110 nm. Alternating laser excitation was provided by a laser combiner (LightHUB Ultra, Omicron) equipped with an acousto-optic tunable filter triggered by the camera digital trigger out via a multifunction I/O device (PCIe-6323, National Instruments). For excitation of AlexaFluor 488, ATTO565 and ATTO647N laser excitation was alternated between 488 nm, 561 nm and 640 nm at intensities on the order of 100 W cm^-2^, 50ms exposure time per excitation wavelength producing 3 camera frames with no time delay between wavelengths. This sequence was repeated every 700 ms to generate a time sequence. Simultaneous imaging of two emission wavelength bands on two halves of the camera sensor was achieved using an image splitter (Optosplit II, Cairn). Two sets of filter combinations (Chroma) were used. For simultaneous imaging of AlexaFluor 488 emission and combined ATTO565 and ATTO647N emission, a ZT543rdc dichroic with ET570LP and ET525/50m emission filters. For simultaneous imaging of ATTO565 emission and ATTO647N emission a ZT633rdc dichroic with ET600/50m and ET700/75m emission filters.

*In situ* on the microscope, both outlets were emptied by aspirating. A 200 µl of a mixture of 1 nM TW and 200 pM DNA track was prepared at room temperature and immediately pipetted into the observation channel outlet at 1 atmosphere. All flows where set to 0.5 µl/min and a rolled kimwipe tissue paper (Kimtech) inserted into the common waste outlet to wick away waste buffer. The surface density of reagents was observed in the observation channel approximately 1 mm away from the intersection. Once single molecule binding on the surface began to reach 1 µm^-2^, as judged by visual inspection, the flows were set to operating values of 7 µl/min (desired solution) and 1 µl/min (two alternative solutions), and the tissue was removed from the waste outlet. An unexposed field of view was selected, and the experiment was started. During the experiment, the solution present in the main observation channel was altered by changing the driving (7 µl/min) buffer every 7 s following the patterns indicated in Figs 4F, 4J, S7A, S8A and S9A. This was achieved in an automated fashion using Fluigent lineup pumps, flow units and OxyGEN software (Fluigent).

### Single molecule FRET analysis

The pixel coordinates of bright spots (Extended Data Fig. 15), independent of time point and color channel, were found using the sci-kit image package^58^ as follows. For each color channel a maximum intensity, *I*, projection over time was calculated. For each of the three color channels this projection was normalized using the following equation:

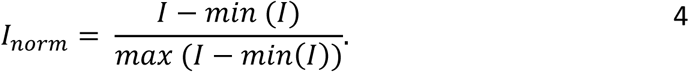

From this normalized maximum intensity projection image, a scale-space volume is generated by computing the Laplacian of Gaussian filtered image with successively increasing standard deviation and generating a stack of images from the result. Single particles were detected using the local maxima in this scale-space volume^59^. Detections close to the edges (35 pixels) of the field of view were removed.

The list of detected coordinate centers was filtered to include only detections that exhibited colocalization between the donor and FRET acceptor channels indicating the presence of both DNA and TW (Extended Data Fig. 15). This was achieved by first identifying nearest neighbor pairs with one member in the donor channel and one in the FRET channel. Then this list of pairs of detections was filtered to only include nearest neighbors closer than 330 nm (3 pixels) of each other. The 330 nm distance threshold was chosen to allow for localization uncertainty, alignment of the optosplit and chromatic aberration.

For each colocalization that was identified between the FRET acceptor and donor channels a background subtracted intensity trace over time was extracted from the full unnormalized image sequence. The background subtracted intensity of the molecule was calculated for each frame by taking the average intensity of the pixels within a radius of 3 pixels (Extended Data Fig. 16A) minus the median of the intensities in the pixels a radius of 4 pixels away (Extended Data Fig. 16B). This was done for each frame and channel, resulting in the traces shown in Extended Data Fig. 17. These traces were matched to the ligand solution in the observation channel at the time of the image acquisition by comparing exact start times and elapsed times from the pump log and metadata of the image. Any trace where the intensity of the first frame was either ∼2.5 times greater than, or ∼0.2 times less than, the average single fluorophore intensity was considered a spurious detection (for example aggregates or unidentified debris) and removed from the analysis. This only resulted in the removal of approximately 10% of colocalizing traces indicating successful preparation of a monodisperse 1:1 ratio of single molecule DNA:TW complexes fixed to the coverslip.

Traces were sorted based on the last detected falling edge in either color channel. Edges (Extended Data Fig. 18A) were determined by Gaussian filtering with a standard deviation of 1.2 (Extended Data Fig. 18B) and taking the first derivative (Extended Data Fig. 18C). Peaks and troughs were identified using sci-pys peakfinder^60^ using a threshold of 2.5 (Extended Data Fig. 18C). Each trace was then normalized between its max value as 1 and the average value of the final 15 frames of the trace as 0. By mapping this normalized data to colors, sorted traces are shown in the kymographs in Figs 4H and L. In Experiment 1, 24-bit RGB color mapping was used with acceptor intensity represented by red and donor intensity by green. In Experiment 2, CMYK color space mapping was used with Atto647N (red) acceptor intensity represented by magenta and Atto565 (yellow) acceptor intensity represented by cyan.

Thresholding of the normalized intensity traces was used to determine the time which TW spent either bound to the intended (adjacent) *trpR* site or bound to the non-adjacent *trpR* site farthest from the bound foot. Due to the differing FRET efficiencies of the two fluorophores, independently selected absolute thresholds were used, specifically 0.8 for red FRET and 0.5 for yellow FRET. Frames in the first buffer condition for which only the red FRET intensity exceeded the specified threshold were counted as intended binding. Frames in the third buffer condition for which only the yellow FRET intensity exceeded the specified threshold were also counted as intended binding. Otherwise, frames were counted as non-adjacent binding including frames in which neither or both FRET intensities exceeded the threshold. To include an equal amount of data from each ligand solution, only the first three solution conditions were included in the analysis and only traces where the protein remained colocalized with the DNA for the full duration of this period were included. The fraction of time spent in the intended binding sites was calculated as 𝑡_𝑖𝑛𝑡𝑒𝑛𝑑𝑒𝑑_/(𝑡_𝑖𝑛𝑡𝑒𝑛𝑑𝑒𝑑_ + 𝑡_𝑢𝑛𝑖𝑛𝑡𝑒𝑛𝑑𝑒𝑑_) for each trace resulting in a mean of 65% ± 7.7% while the fraction of time spent in the unintended binding sites was calculated as 𝑡_𝑢𝑛𝑖𝑛𝑡𝑒𝑛𝑑𝑒𝑑_/(𝑡_𝑖𝑛𝑡𝑒𝑛𝑑𝑒𝑑_ + 𝑡_𝑢𝑛𝑖𝑛𝑡𝑒𝑛𝑑𝑒𝑑_) for each trace resulting in a mean of 35% ± 6.3% where limits are 95% confidence intervals.

## Acknowledgments

This project has evolved over two decades with input from many scientists. We acknowledge the contributions made by group members: Martin Zuckermann, Olivier Laprévote, Nathan Kuwada, Ben Lopez, Aimee Boyle, Drew Thompson, Laleh Samii, Suzana Kovacic, Hasti Iranmanesh, James Walsh, Christopher Angstmann, Jowa Chan, Richard Sessions, Marc Bruning, Lara Small, Ben Clifton, Martina Balaz and Michael Konopik. We also thank Nicholas Dixon, Lisanne Spenkelink, Daniela Stock, Jonas Tegenfeldt, Lawrence Lee, Sophie Hertel and Slobodan Jergic for advice and guidance at various stages of this project.

The authors acknowledge the use of facilities in the Structural Biology Facility within the Mark Wainwright Analytical Centre-UNSW. We thank UNSW Single Molecule Science for generous access to SPR facilities.

## Funding

Human Frontiers in Science Program grant RGP0031/2007 (HL, PMGC, NRF, DNW)

European Union’s Horizon 2020 research and innovation programme under grant agreement No 951375 (ArtMotor) (HL, BH, PMGC)

Australian Research Council Discovery Project grant DP170103153 (PMGC, TB)

Natural Sciences and Engineering Research Council of Canada (NSERC) grants RGPIN-2015-05545, RGPIN-2020-04680 (NRF)

Swedish Research Council project numbers 2015-03824 (HL), 2020-04226 (HL), 2020-05266 (RE), 638-2013-9243 (RE).

## Author contributions

Conceptualization: RBD, DNW, NRF, HL, PMGC

Methodology: PN, NOR, NG, RBD, CWL, AL, RE, CSN, GAB, EHCB, AEW, APD, JPB, PJ, TB, BH, DNW, NRF, HL, PMGC

Investigation: PN, NOR, NG, RBD, CWL, AL, RE, AEW, APD, BH, DNW, NRF, HL, PMGC Visualization: PN, NOR, NG, RBD, CWL, AL, RE, AEW, APD, NRF, HL, PMGC

Funding acquisition: RE, TB, BH, DNW, NRF, HL, PMGC Project administration: BH, DNW, NRF, HL, PMGC Supervision: BH, DNW, NRF, HL, PMGC

Writing – original draft: PN, NOR, NG, CWL, HL, PMGC

Writing – review & editing: PN, NOR, NG, RBD, CWL, AL, RE, CSN, GAB, EHCB, AEW, APD, INU, JPB, PJ, TB, BH, DNW, NRF, HL, PMGC

## Competing interests

Authors declare that they have no competing interests.

## Materials & Correspondence

Correspondence and material requests should be addressed to PMGC and HL.

## Data availability

SAXS data have been deposited with the Small Angle Scattering Biological Data Bank (SASBDB) with the accession code SASDWM8. All other data are available in the main text or the supplementary materials.

**Supplementary Information** is available for this paper.

## Extended Data Table/Figure Legends

**Extended Data Table 1.**
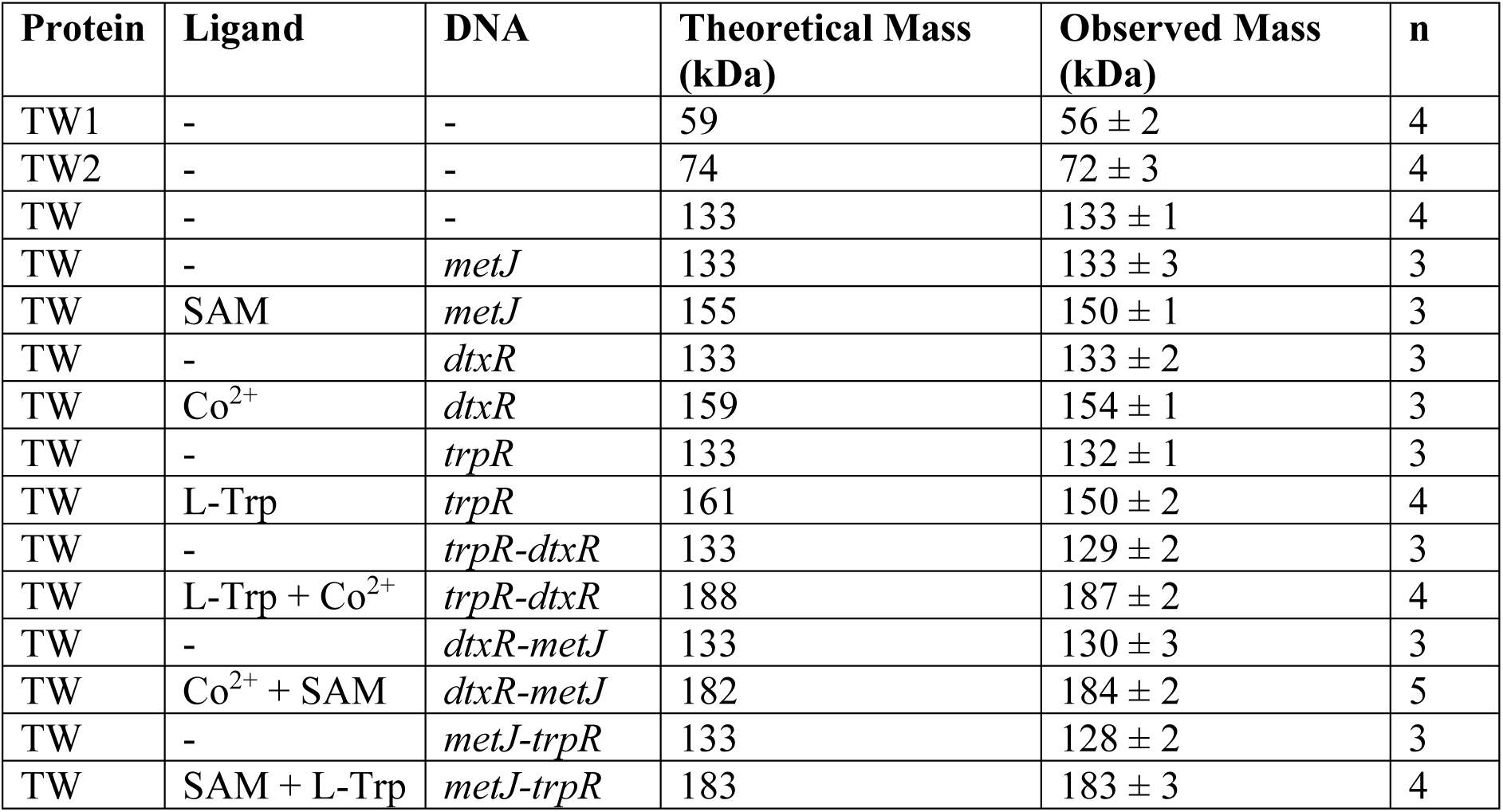
TW construct and complex molecular masses. Protein and protein:DNA molecular masses were determined using mass photometry, as shown in Fig. 3C & Extended Data Fig. 5. Theoretical masses were calculated assuming TW and TW constructs act as obligate dimers. Observed masses represent average ± SEM, calculated from replicate experiments with sample size of n.

| Protein | Ligand | DNA | Theoretical Mass (kDa) | Observed Mass (kDa) | n |
| --- | --- | --- | --- | --- | --- |
| TW1 | - | - | 59 | 56 ± 2 | 4 |
| TW2 | - | - | 74 | 72 ± 3 | 4 |
| TW | - | - | 133 | 133 ± 1 | 4 |
| TW | - | *metJ* | 133 | 133 ± 3 | 3 |
| TW | SAM | *metJ* | 155 | 150 ± 1 | 3 |
| TW | - | *dtxR* | 133 | 133 ± 2 | 3 |
| TW | Co<sup>2+</sup> | *dtxR* | 159 | 154 ± 1 | 3 |
| TW | - | *trpR* | 133 | 132 ± 1 | 3 |
| TW | L-Trp | *trpR* | 161 | 150 ± 2 | 4 |
| TW | - | *trpR-dtxR* | 133 | 129 ± 2 | 3 |
| TW | L-Trp + Co<sup>2+</sup> | *trpR-dtxR* | 188 | 187 ± 2 | 4 |
| TW | - | *dtxR-metJ* | 133 | 130 ± 3 | 3 |
| TW | Co<sup>2+</sup> + SAM | *dtxR-metJ* | 182 | 184 ± 2 | 5 |
| TW | - | *metJ-trpR* | 133 | 128 ± 2 | 3 |
| TW | SAM + L-Trp | *metJ-trpR* | 183 | 183 ± 3 | 4 |

**Extended Data Table 2.** TW:DNA affinities in the presence of a single ligand by SPR. Affinities were determined by titrating TW onto a surface containing DNA. The average steady state response units in the contact phase were fitted to the Hill equation to determine the apparent *K_d_* and Hill coefficient (h). Values represent average ± SEM, calculated from replicate experiments with sample size of n.

| DNA | Protein | Ligand | Apparent $K_d$ (nM) | h | n |
| --- | --- | --- | --- | --- | --- |
| *metJ* | MetJ | SAM | $4 \pm 0.4$ | $1.22 \pm 0.02$ | 4 |
| *metJ* | TW | SAM | $18 \pm 5$ | $0.96 \pm 0.07$ | 4 |
| *dtxR* | DtxR | $\text{Co}^{2+}$ | $5.5 \pm 0.2$ | $0.99 \pm 0.03$ | 5 |
| *dtxR* | TW | $\text{Co}^{2+}$ | $8 \pm 1$ | $0.92 \pm 0.10$ | 4 |
| *trpR* | TrpR | L-Trp | $48 \pm 2$ | $1.61 \pm 0.02$ | 4 |
| *trpR* | TW | L-Trp | $27 \pm 4$ | $1.07 \pm 0.03$ | 4 |
| *trpR-dtxR* | TW | $\text{Co}^{2+}$ | $3 \pm 2$ | $1.0 \pm 0.2$ | 2 |
| *trpR-dtxR* | TW | L-Trp | $15 \pm 4$ | $1.27 \pm 0.09$ | 4 |
| *dtxR-metJ* | TW | SAM | $14 \pm 2$ | $1.2 \pm 0.2$ | 3 |
| *dtxR-metJ* | TW | $\text{Co}^{2+}$ | $5 \pm 2$ | $1.2 \pm 0.1$ | 3 |
| *dtxR-metJ* | TW | L-Trp | $50 \pm 20$ | $1.72 \pm 0.03$ | 3 |
| *metJ-trpR* | TW | SAM | $18 \pm 2$ | $1.3 \pm 0.2$ | 3 |
| *metJ-trpR* | TW | L-Trp | $13 \pm 4$ | $1.22 \pm 0.09$ | 4 |

**Extended Data Table 3.** TW:DNA dissociation kinetics by SPR. TW was loaded onto a DNA surface in the presence of the relevant ligands. Dissociation was triggered by injecting a solution containing no TW, the relevant ligands, and 1 µM DNA. 1 µM of each of the relevant single site DNA oligonucleotides were used for double ligand dissociation experiments. Dissociation curves were fitted to a single exponential decay function to determine apparent k_off_. Half-lives were calculated from the apparent k_off_’s. Values represent average ± SEM, calculated from replicate experiments with sample size of n.

| DNA | Ligand | Apparent $k_{\text{off}}$ ($\text{s}^{-1}$) | Apparent Half-life (s) | n |
| --- | --- | --- | --- | --- |
| *metJ* | No ligand | $3.7 \pm 0.3$ | $0.19 \pm 0.02$ | 3 |
| *metJ* | SAM | $1.14 \pm 0.06$ | $0.61 \pm 0.03$ | 3 |
| *dtxR* | No ligand | $0.45 \pm 0.02$ | $1.56 \pm 0.07$ | 3 |
| *dtxR* | $\text{Co}^{2+}$ | $0.164 \pm 0.009$ | $4.2 \pm 0.4$ | 3 |
| *trpR* | No ligand | $0.99 \pm 0.02$ | $0.70 \pm 0.02$ | 3 |
| *trpR* | L-Trp | $0.36 \pm 0.04$ | $2.0 \pm 0.2$ | 3 |
| *trpR-dtxR* | L-Trp + $\text{Co}^{2+}$ | $0.0239 \pm 0.0008$ | $29 \pm 0.6$ | 3 |
| *dtxR-metJ* | SAM + $\text{Co}^{2+}$ | $0.0145 \pm 0.0005$ | $48 \pm 1$ | 2 |
| *metJ-trpR* | SAM + L-Trp | $0.0233 \pm 0.002$ | $29.7 \pm 0.2$ | 3 |

**Extended Data Table 4.**
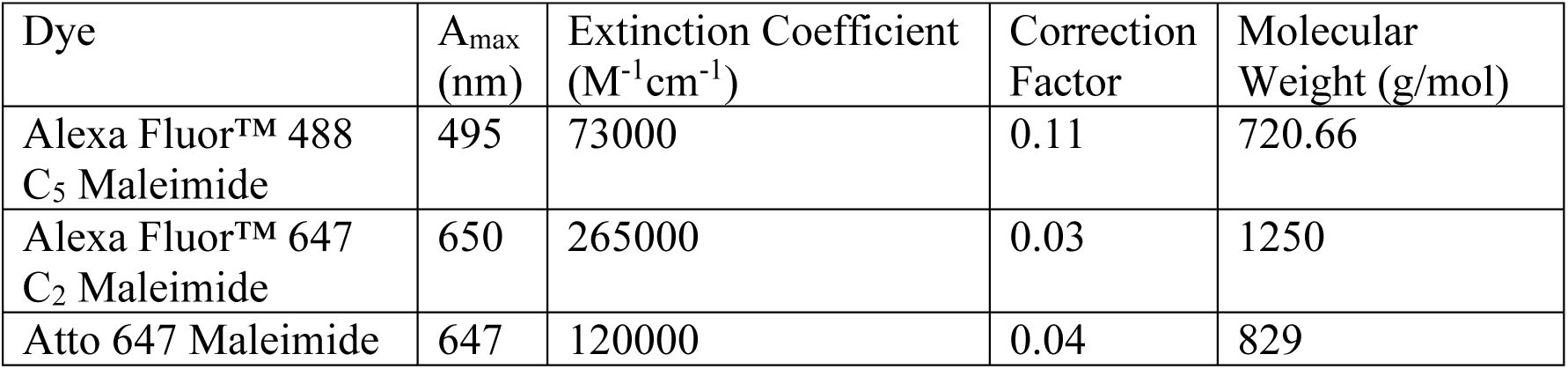
Properties of dye conjugates used for protein labelling. Data from Thermofisher Scientific and Sigma Aldrich.

**Extended Data Table 5.**
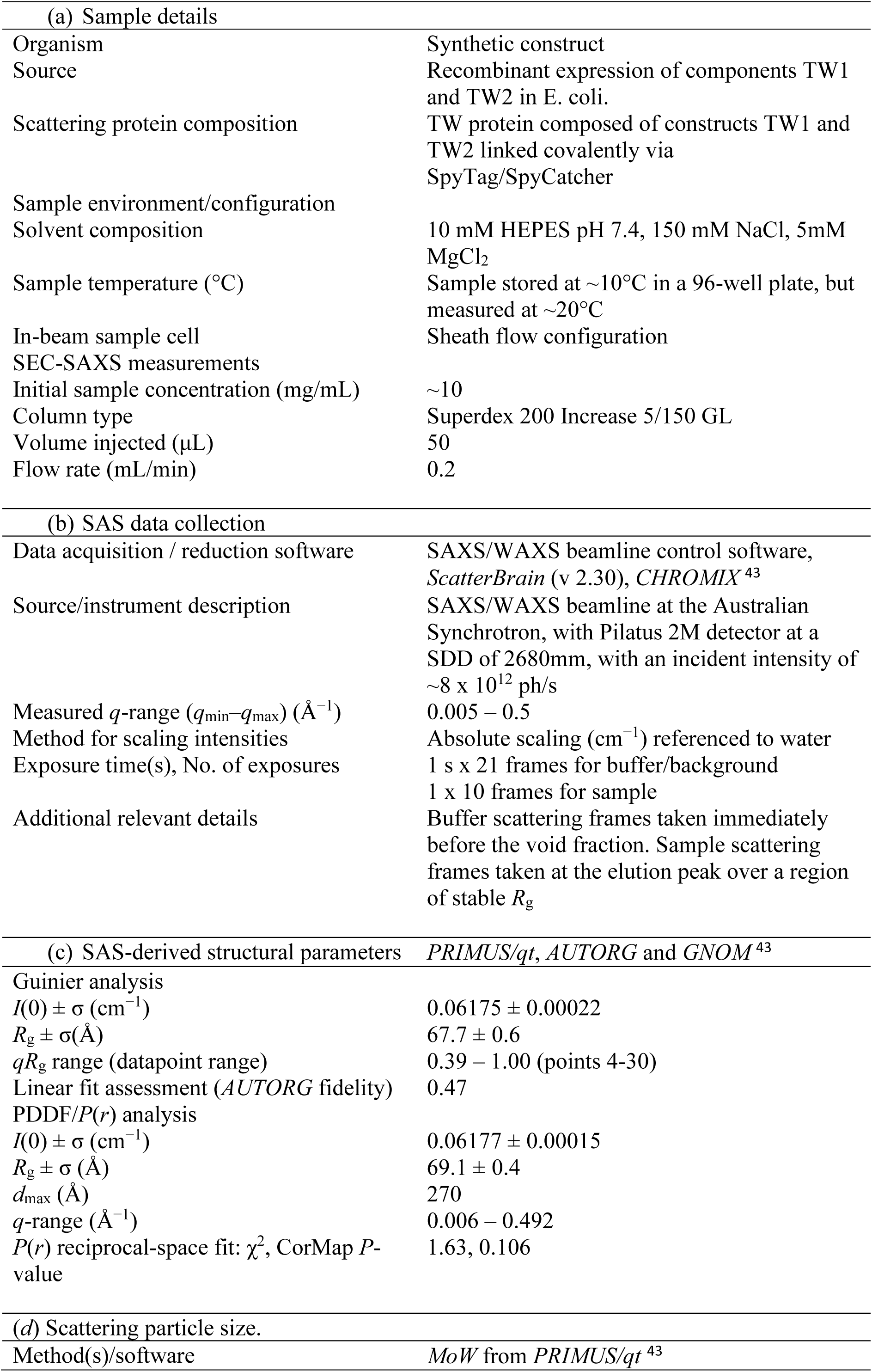
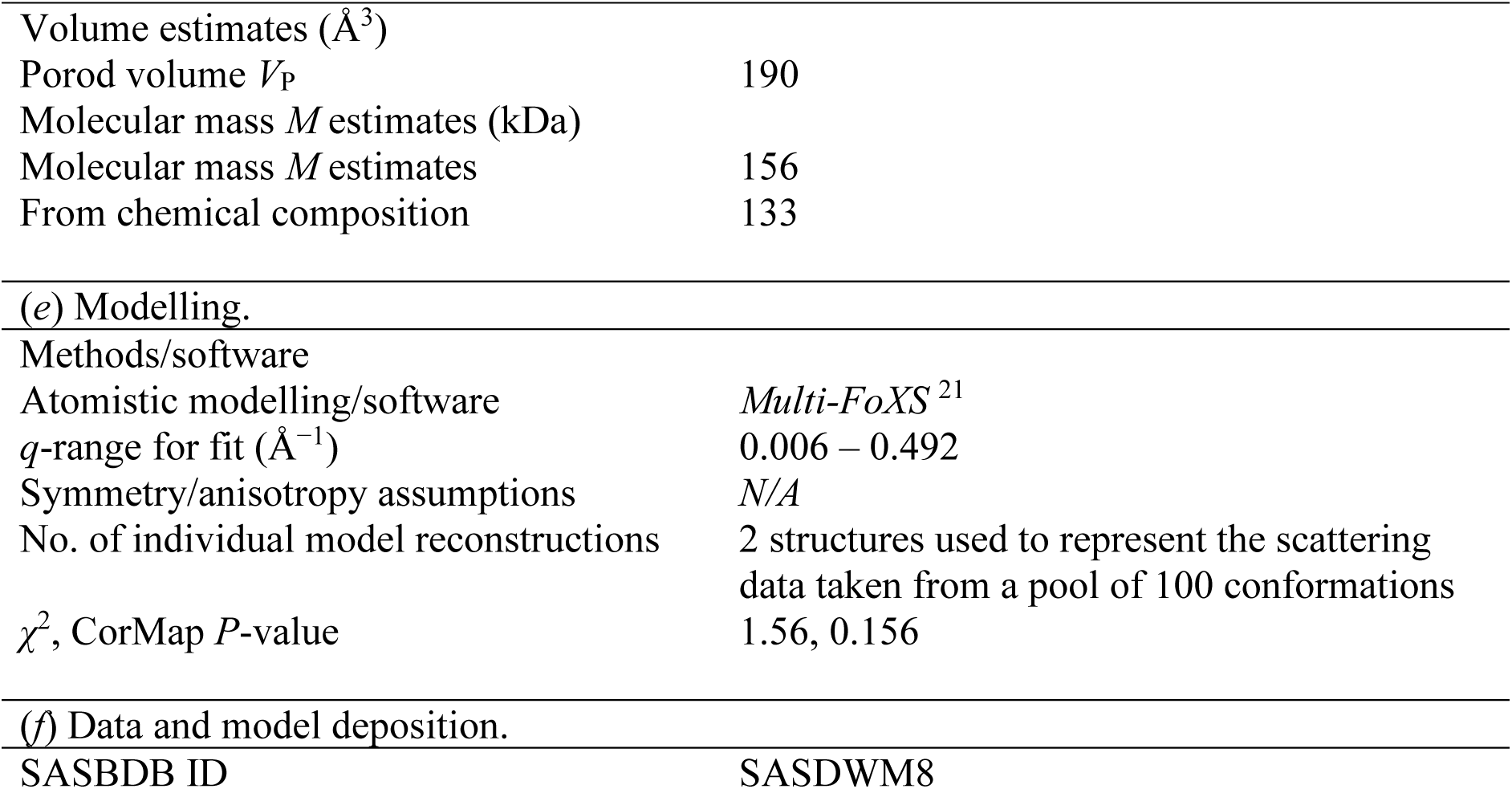
SEC-SAXS data collection, reduction and analysis.

**Extended Data Table 6.**
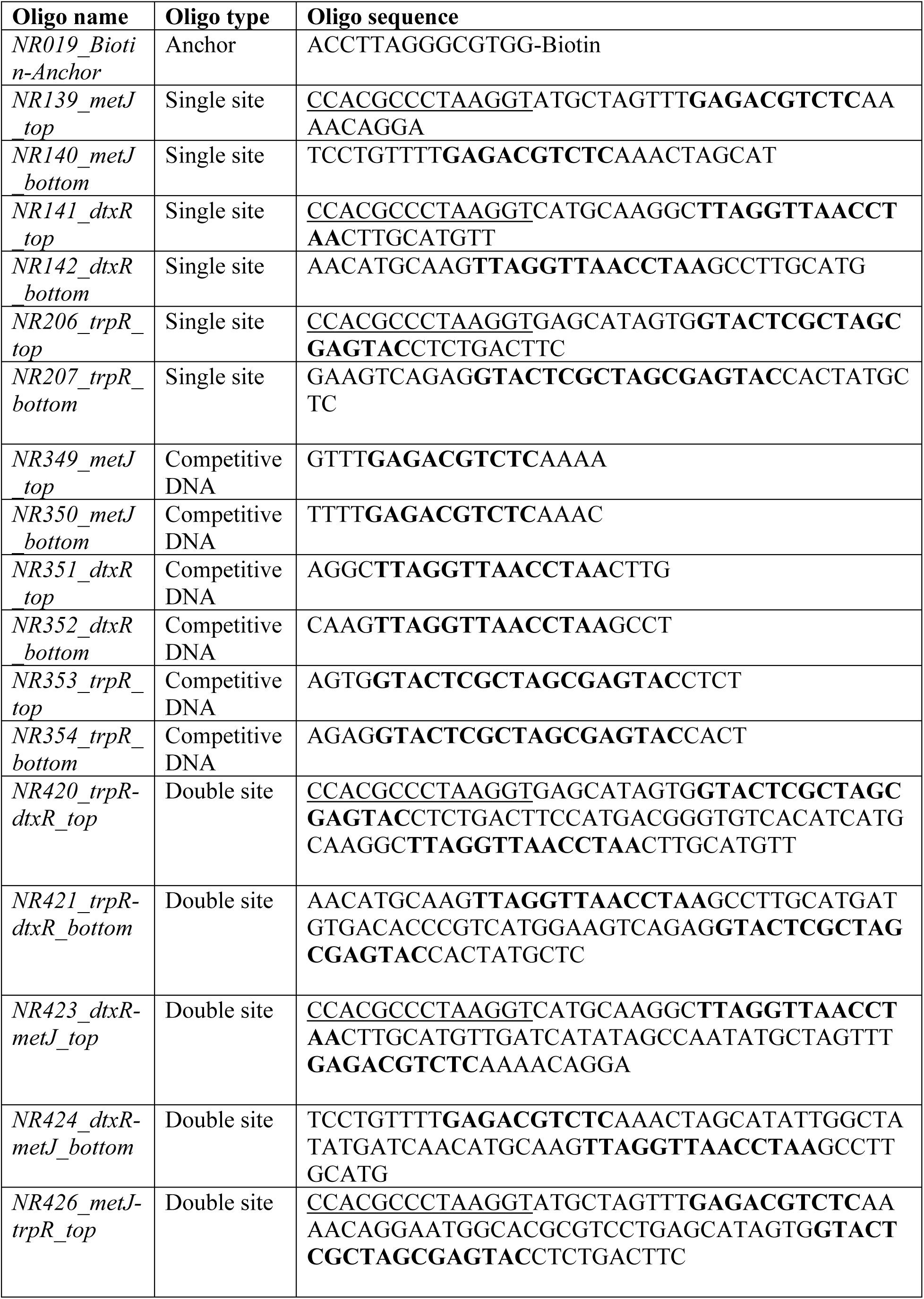
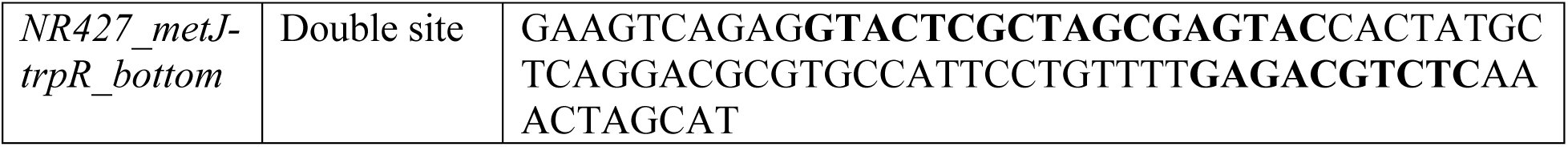
Oligonucleotide sequences used for SPR and mass photometry. Underlined sequences represent complements to Biotin-Anchor strand used to anneal DNA surfaces in SPR experiments. Single- and double-site oligos refer to sequences that contain one and two repressor cognate sites, respectively. Competing DNA refers to sequences containing a single cognate binding site that was used to trigger dissociation in SPR experiments. Bold sequences represent the cognate binding sequences for the relevant repressor. Top and bottom sequences were annealed to form double-stranded DNA for use in experiments.

**Extended Data Table 7.**
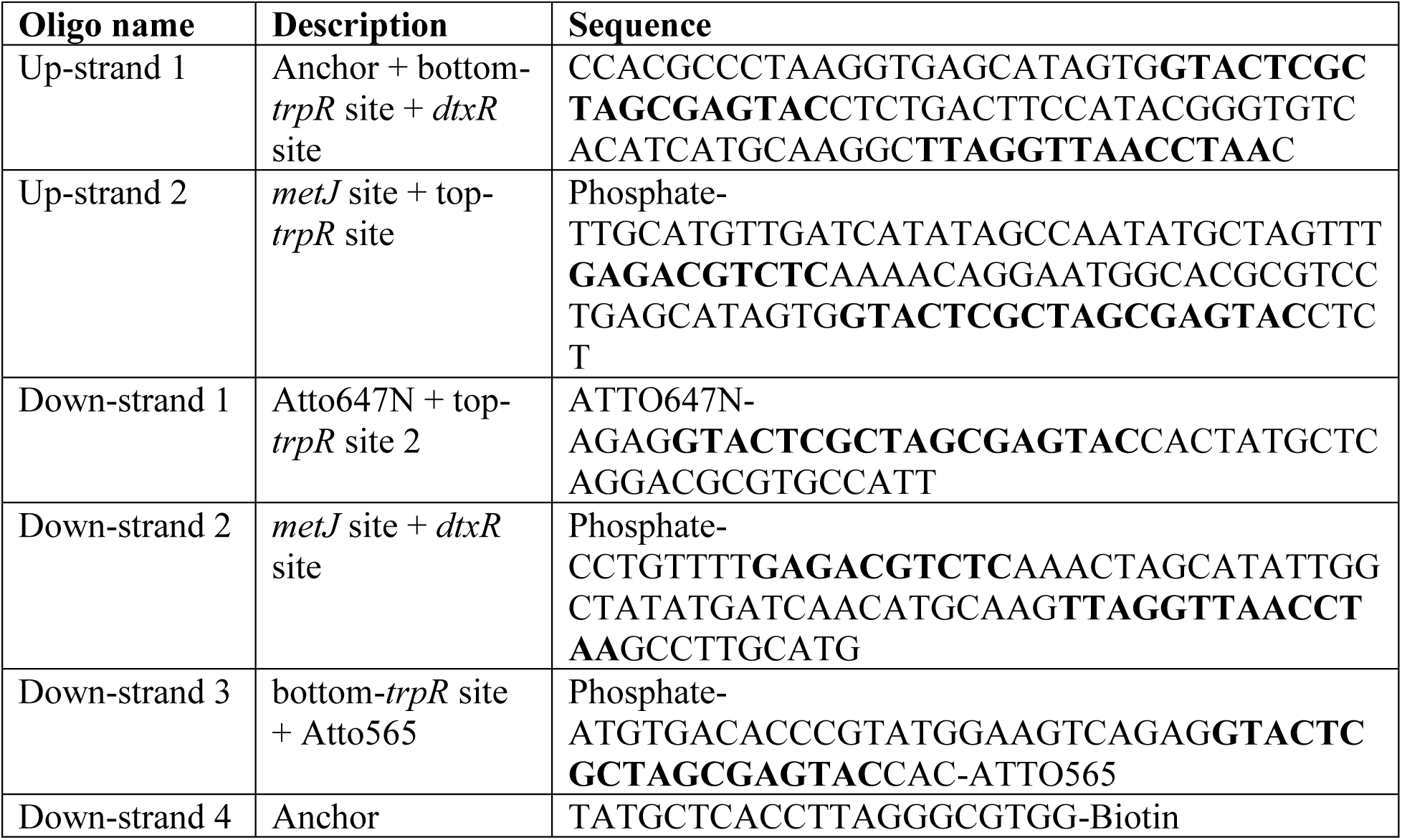
Oligonucleotide sequences used to construct the track for single molecule FRET. The four-site track was annealed, ligated, and purified as described in Materials and Methods from the six oligonucleotides listed here. Up-strands refer to strands designed with 5’ to 3’ direction away from the coverslip. Down-strands refer to strands designed with 5’ to 3’ direction towards the coverslip. Bold sequences represent the cognate binding sequences for the relevant repressor.

**Extended Data Table 8:** Single- and no-ligand rates used in the simulations for the transition between the unbound and the intended bound state. Single- and no-ligand off-rates are identical to the rates measured in the SPR dissociation experiments (Extended Data Table 3). The on-rates are assumed to be repressor-independent and are chosen to reproduce the measured double-ligand rates as well as possible; the double-ligand rates resulting from the model are all within a factor 3 from the measured rates, compare Extended Data Table 3. They are extracted from the model by simulating the detachment of TW from the track in the presence of fixed ligand combinations Trp + Co^2+^, Co^2+^ + SAM, and SAM + Trp, respectively. Transitions between unbound state and overstepped bound state use the same off-rates, but all on-rates are rescaled by a factor a.

| DNA | Ligand | Off rate (s<sup>-1</sup>) | On rate (s<sup>-1</sup>) |
| --- | --- | --- | --- |
| *metJ* | SAM | 1.14 | 10 |
|  | none | 3.7 | 0 |
| *dtxR* | Co<sup>2+</sup> | 0.164 | 10 |
|  | none | 0.45 | 0 |
| *trpR* | Trp | 0.36 | 10 |
|  | none | 0.99 | 0 |
| *trpR-dtxR* | Trp + Co<sup>2+</sup> | 0.011 | n/a |
| *dtxR-metJ* | SAM + Co<sup>2+</sup> | 0.031 | n/a |
| *metJ-trpR* | SAM + Trp | 0.067 | n/a |

**Extended Data Fig. 1.**
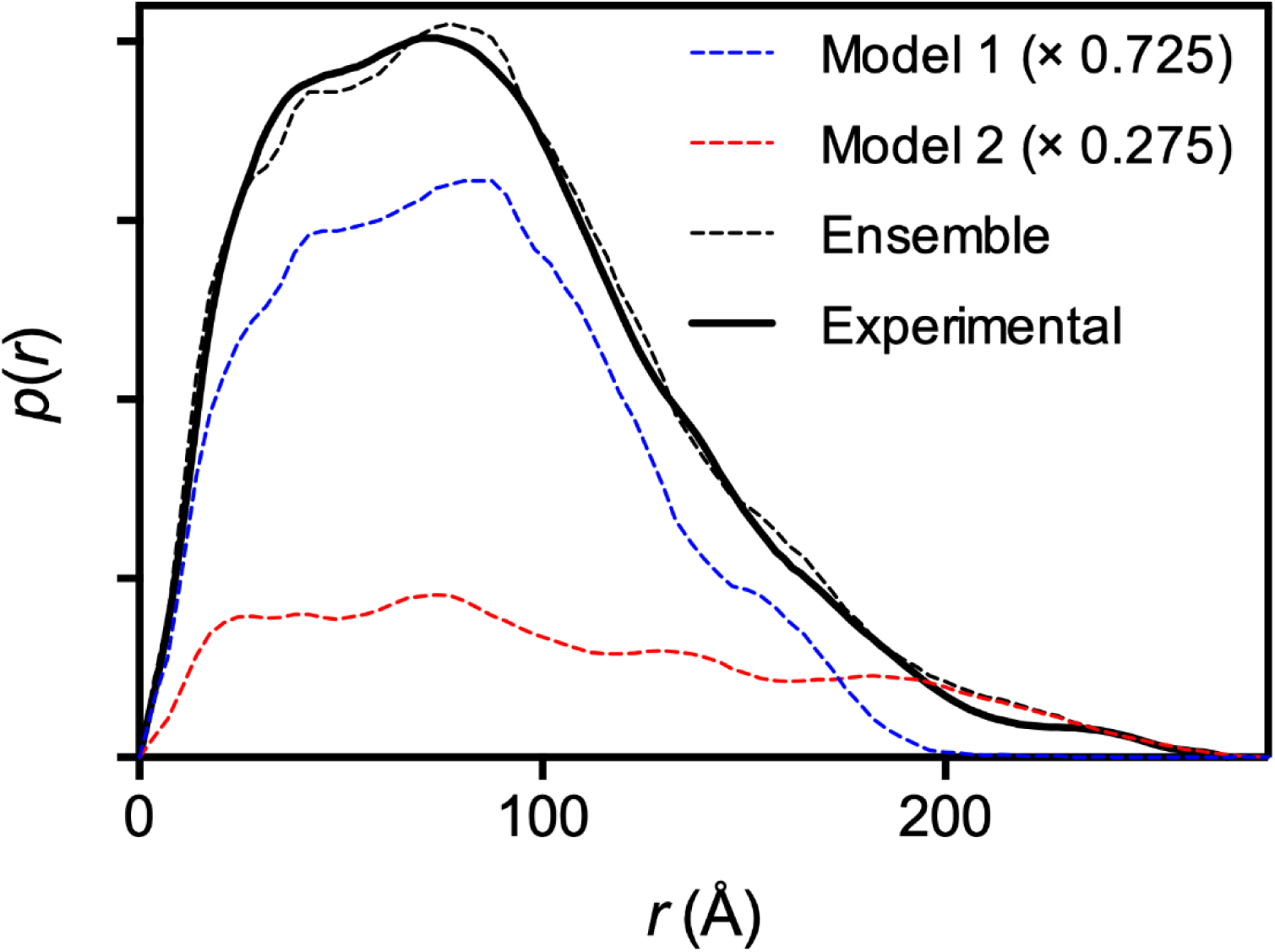
P(r) plot calculated from the SEC-SAXS data. The figure shows the pair-distance distribution function, *P*(*r*), derived from the experimental data (solid black line). Overlaid are the *P*(*r*) functions for the representative compact (dotted blue line) and extended model structures (dotted red line) and their weighted sum representing the ensemble average (black dotted line).

**Extended Data Fig. 2.**
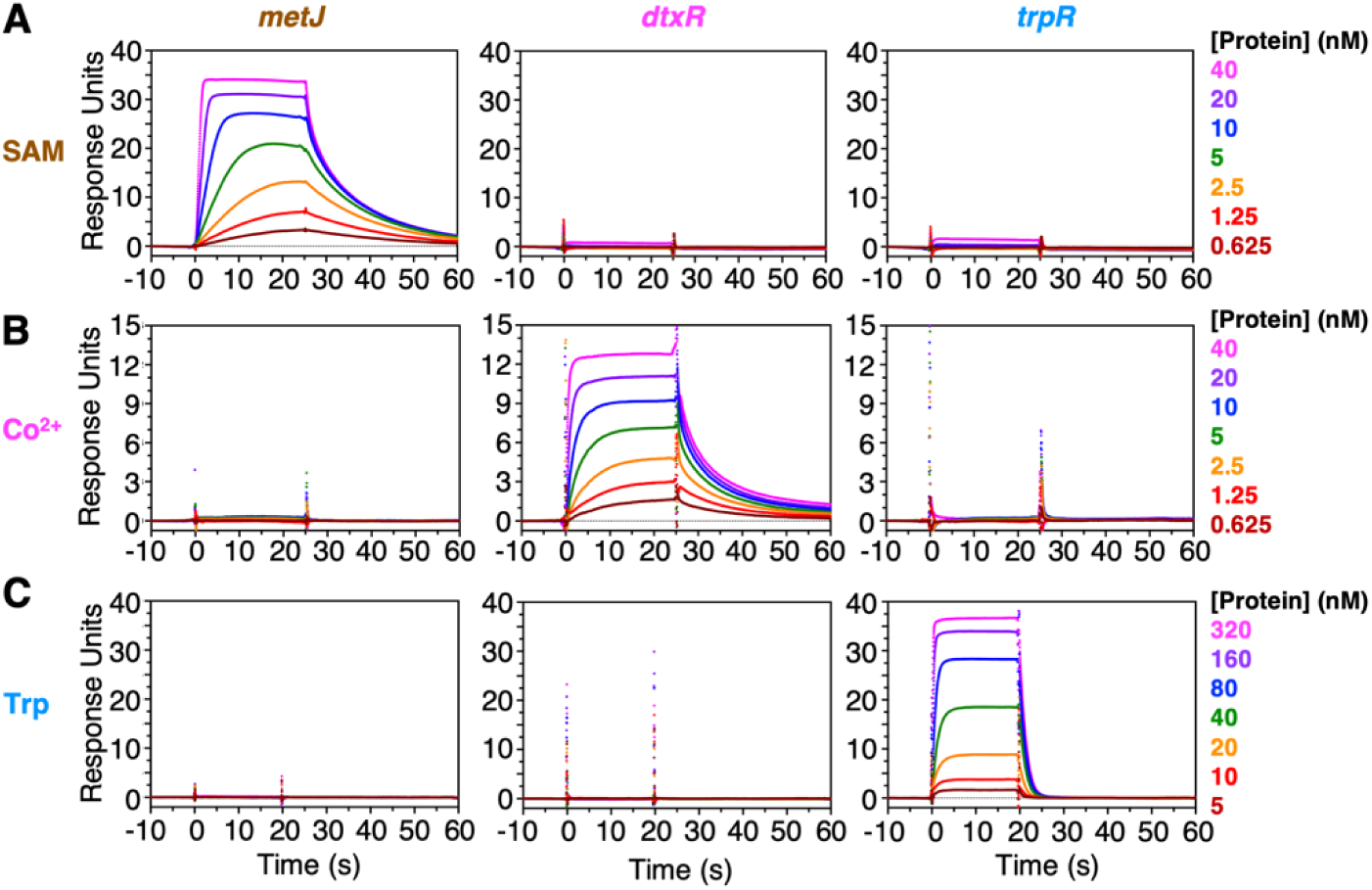
DNA binding by MetJ, DtxR and TrpR is orthogonal. MetJ, DtxR and TrpR were titrated onto each of their respective cognate sites via SPR in the presence of **(A)** SAM, **(B)** Co^2+^ and **(C)** Trp. The DNA sites used in these experiments are named above the figures, with sequences reported in Extended Data Table 6. The steady state responses were fit to the Hill equation, with the fitted binding parameters for each specific interaction reported in Extended Data Table 2.

**Extended Data Fig. 3.**
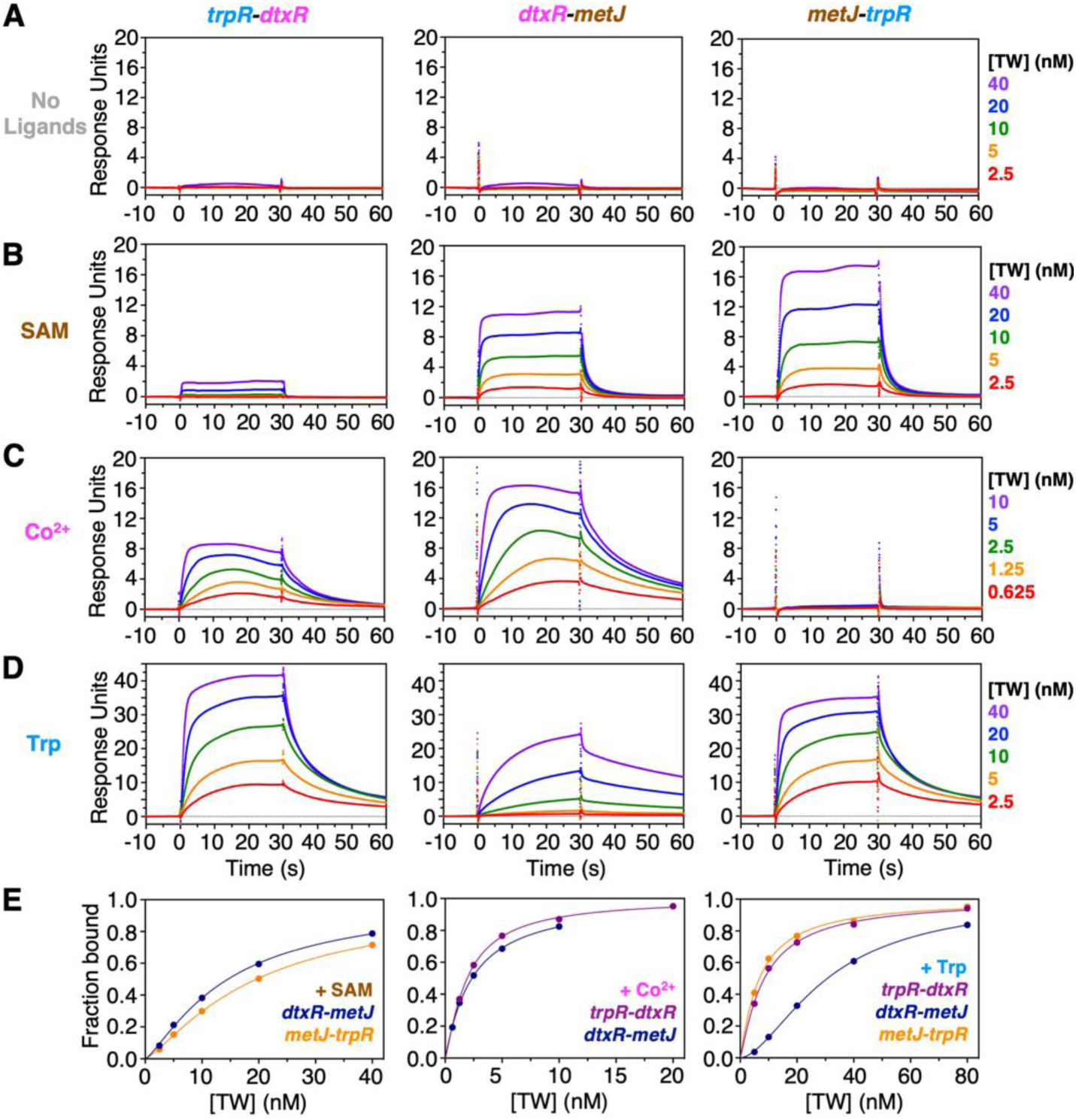
TW binding double-site DNA in the presence of single ligands, by SPR. TW was titrated onto DNA containing two repressor cognate sites **(A)** in the absence of ligand, and in the presence of **(B)** SAM, **(C)** Co^2+^ or **(D)** Trp. The DNA sites used in these experiments are named above the figures, with sequences reported in Extended Data Table 6. **(E)** Representative steady state SPR binding isotherms of TW binding to double-site DNA were fitted to the Hill equation. The ligands and DNA used in these experiments are as per the labels within the figure panels. The fitted parameters for TW binding to the double site DNA are reported in Extended Data Table 2.

**Extended Data Fig. 4.**
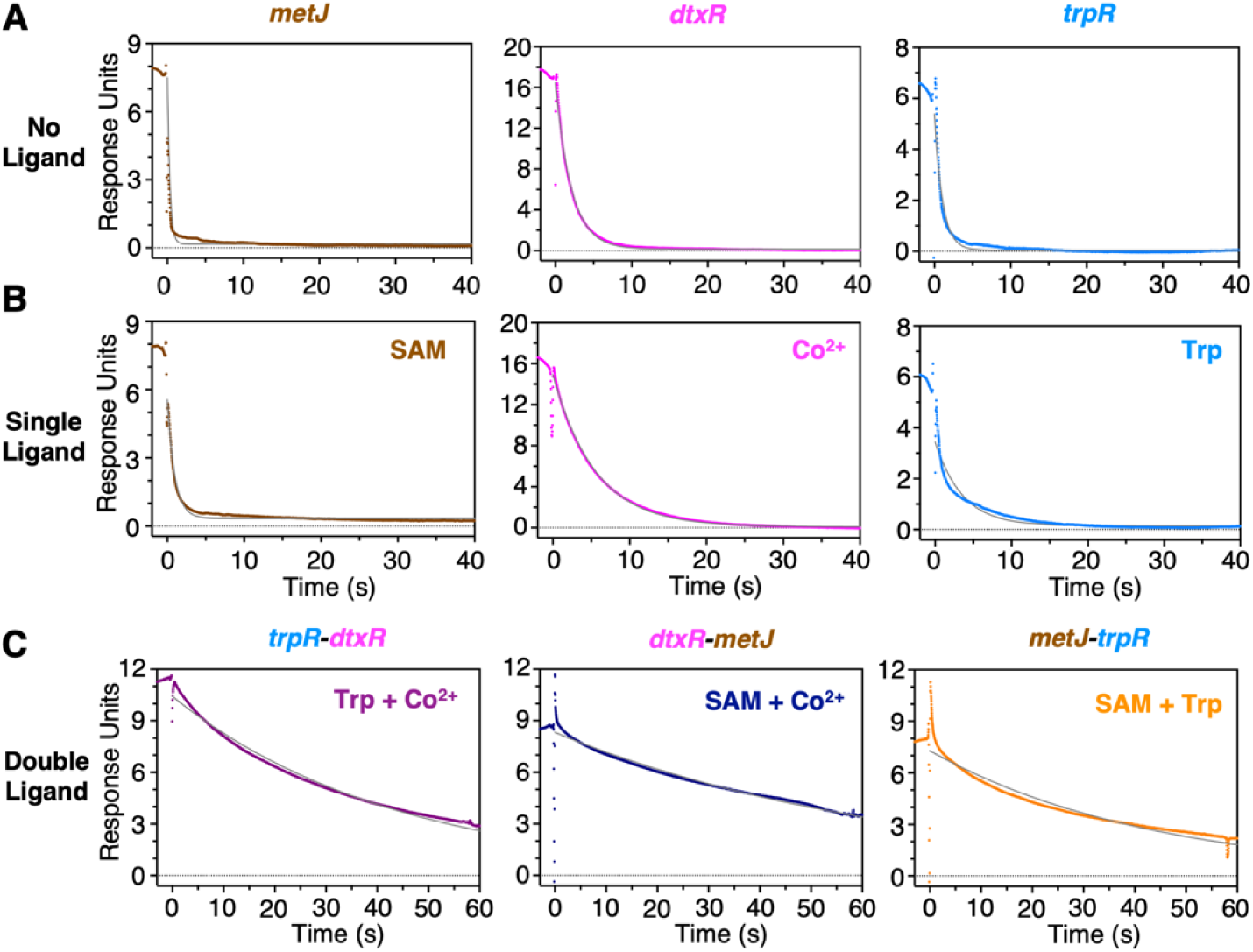
Dissociation kinetics of TW, by SPR. Representative dissociation experiments in which TW was loaded with the relevant ligand onto DNA containing either one or two repressor cognate sites. Dissociation was triggered by injection of a solution containing 1 µM free DNA containing the same cognate site as the DNA bound to the SPR chip surface. Free DNA sequences lacked an SPR anchor overhang and therefore could not bind to the SPR surface. The DNA sites used in these experiments are named above the figures, with sequences reported in Extended Data Table 6. Ligands used in each experiment are labelled within the figure. Curves were fit to a single exponential decay function, with the apparent off-rates reported in Extended Data Table 3. **(A)** Dissociation in the presence of DNA but absence of ligand. **(B)** Dissociation in the presence of both DNA and single ligands. **(C)** Dissociation in the presence of double ligands and two free oligonucleotides, each containing a single relevant repressor cognate site.

**Extended Data Fig. 5.**
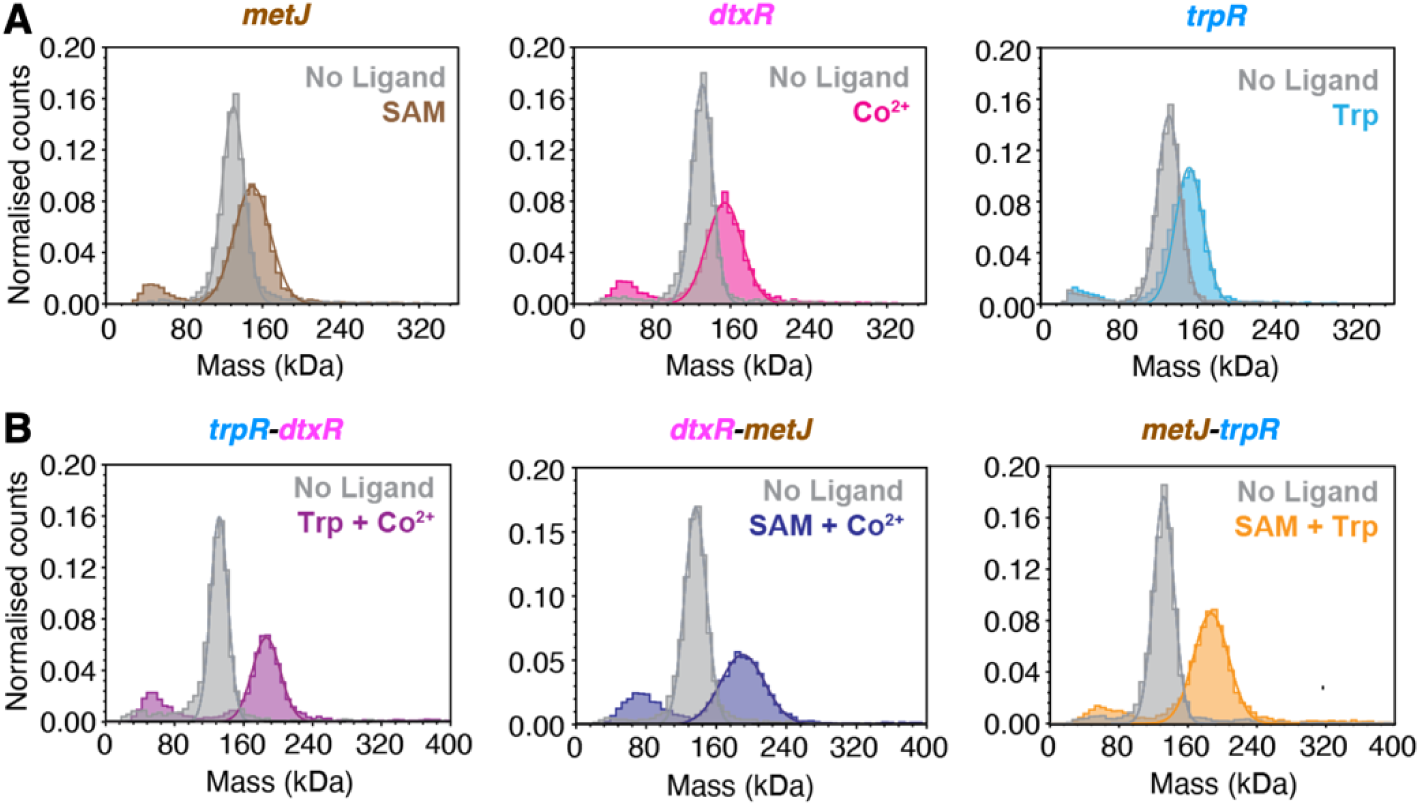
TW forms monodisperse 1:1 complexes with both single- and double-site DNA, by mass photometry. TW:DNA mixtures were subjected to mass photometry. The resulting main mass distributions were fit to a Gaussian function, with molecular masses for each complex reported in Extended Data Table 1. Signal interference by SAM and the binding of DNA to the glass slide in the presence of Co^2+^ resulted in additional mass peaks centered around 60–80 kDa. The DNA sites used in these experiments are named above the figures, with sequences reported in Extended Data Table 6. The ligands used in each experiment are as per the labels within the figure panels. **(A)** TW binding to single-site DNA in the absence and presence of single ligands. The observed molecular masses of the TW:single-site complexes (150–154 kDa) were in good agreement with the theoretical molecular masses (155–161 kDa). **(B)** TW binding to double-site DNA in the absence and presence of double ligands. The observed molecular masses of the TW:double-site complexes (183–187 kDa) were in excellent agreement with the theoretical molecular masses (182–188 kDa).

**Extended Data Fig. 6.**
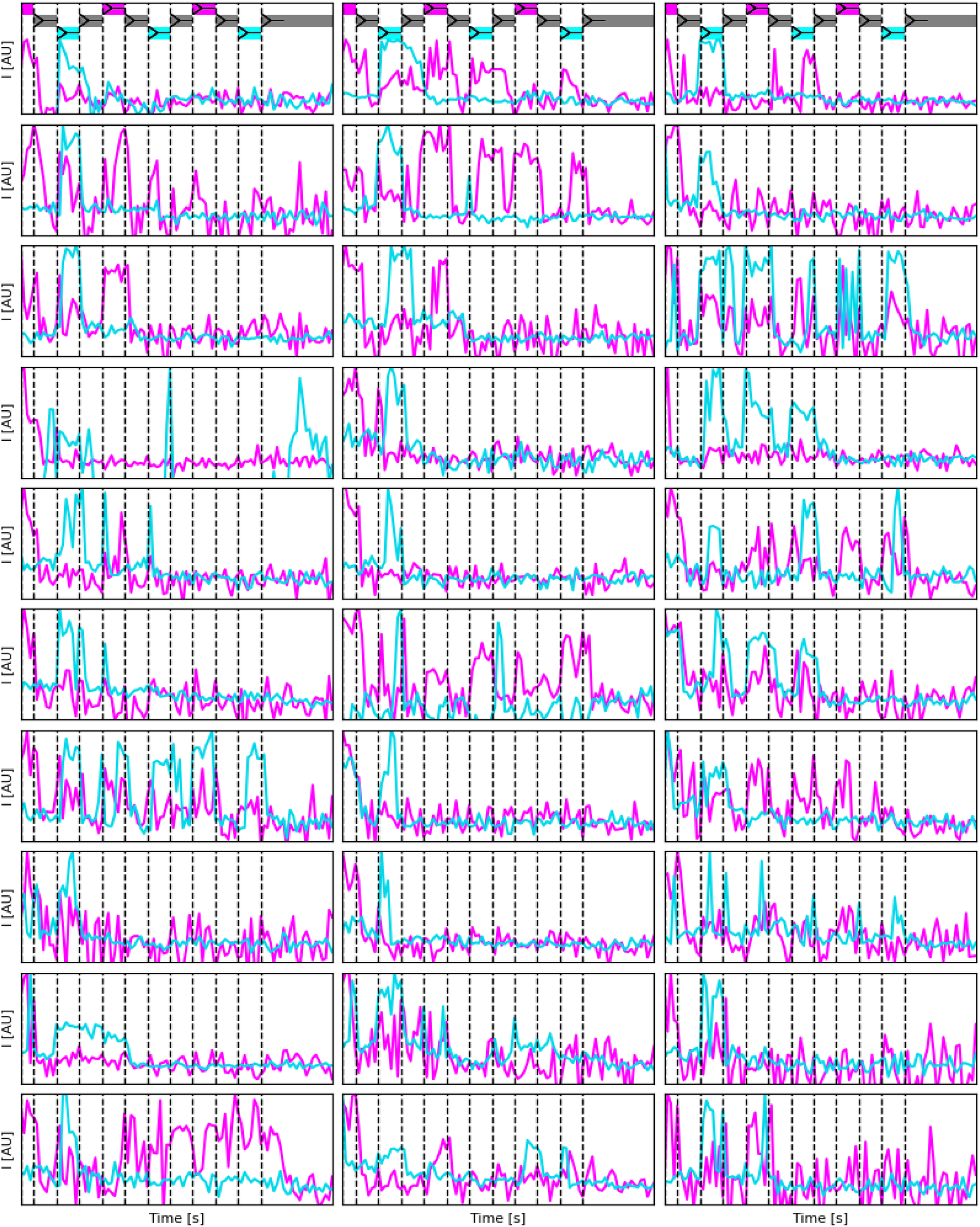
Normalized examples of single-molecule traces using two-colour imaging with emission split at 633 nm to simultaneously record ATTO565 emission (FRET emission from the bottom *trpR* site, shown in cyan) and ATTO647N emission (FRET emission from the top *trpR* site, shown in magenta). At the top of each column, a schematic of expected TW position and false colour representation of expected fluorescence signal given solution changes (dashed lines).

**Extended Data Fig. 7.**
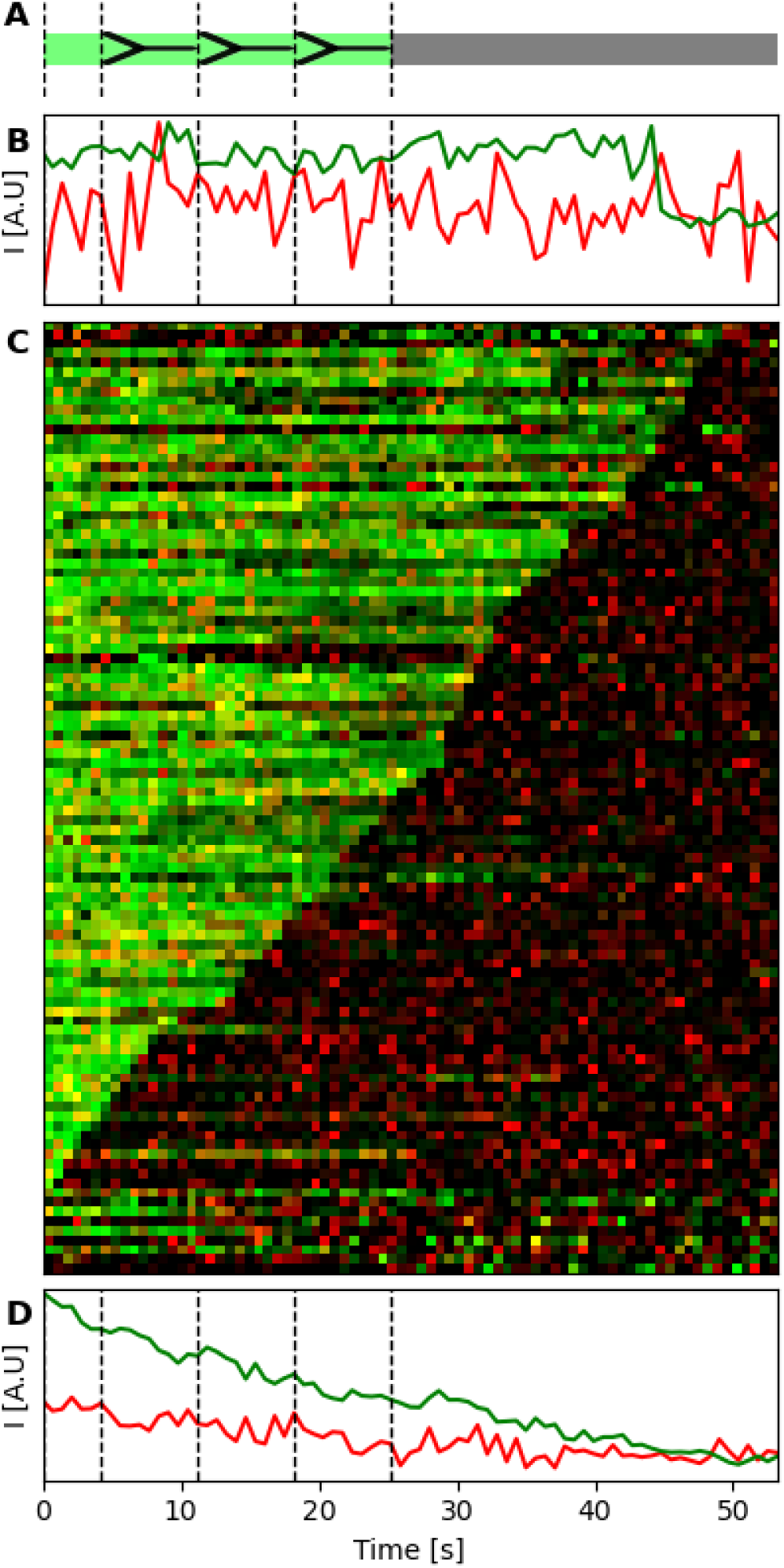
TW remained static on DNA track when ligand combination (Co_2+_+SAM) remained the same after microfluidic solution changes, as observed by single-molecule FRET. **(A)** Schematic of expected TW position and false-color representation of expected fluorescence signal given solution changes (dashed lines). **(B)** Normalized example single-molecule trace. **(C)** Kymographs of all detected colocalizing single molecules sorted by time of last identified fluorescence. **(D)** Sum of all single-molecule traces in the experiment.

**Extended Data Fig. 8.**
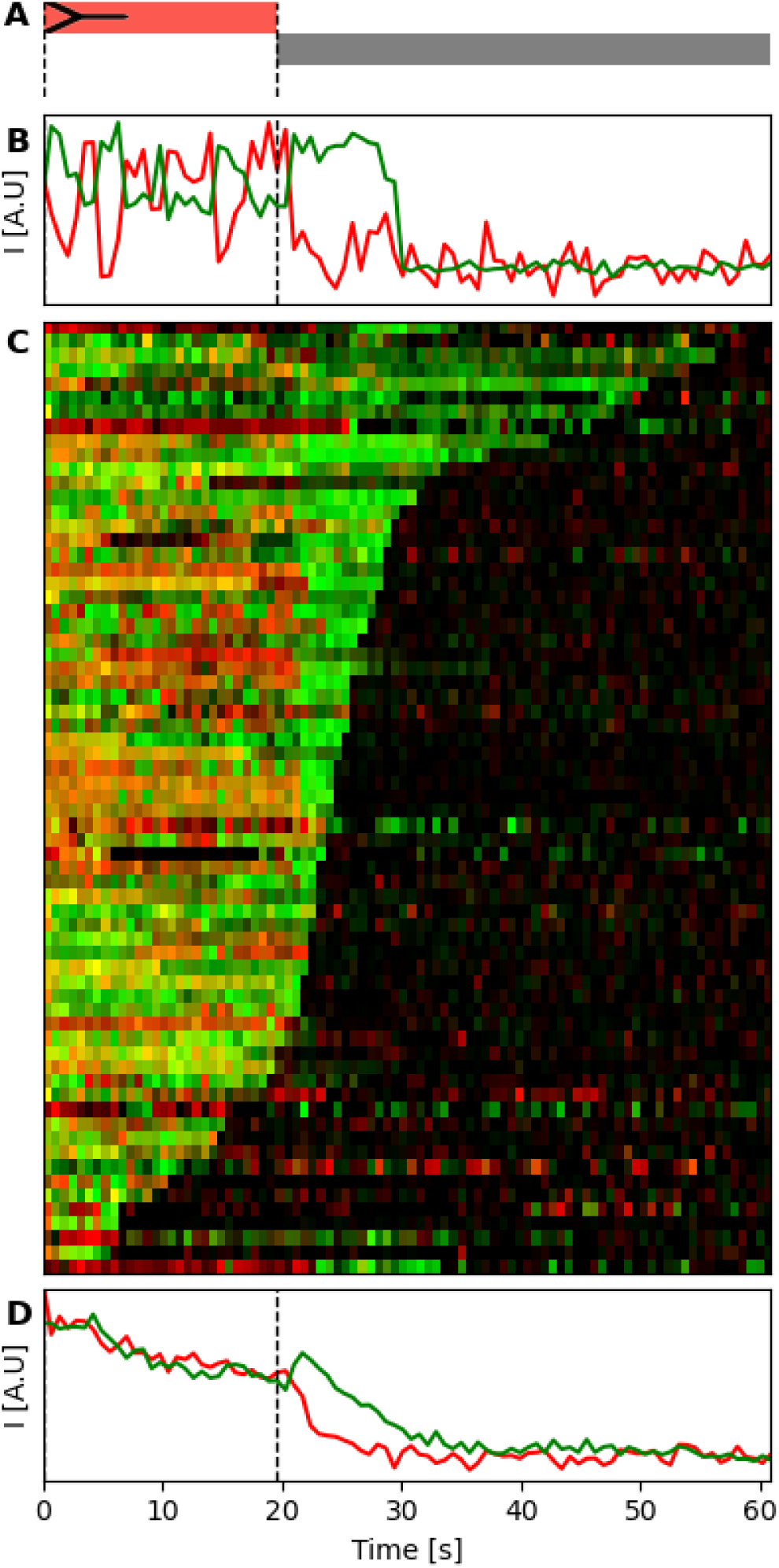
TW rapidly detached from DNA track after removal of ligands (SAM+Trp), as observed by single-molecule FRET. **(A)** Schematic of expected TW position and false-color representation of expected fluorescence signal given solution changes (dashed lines). **(B)** Normalized example single-molecule trace. **(C)** Kymographs of all detected colocalizing single molecules sorted by time of last identified fluorescence. **(D)** Sum of all single-molecule traces in the experiment.

**Extended Data Fig. 9.**
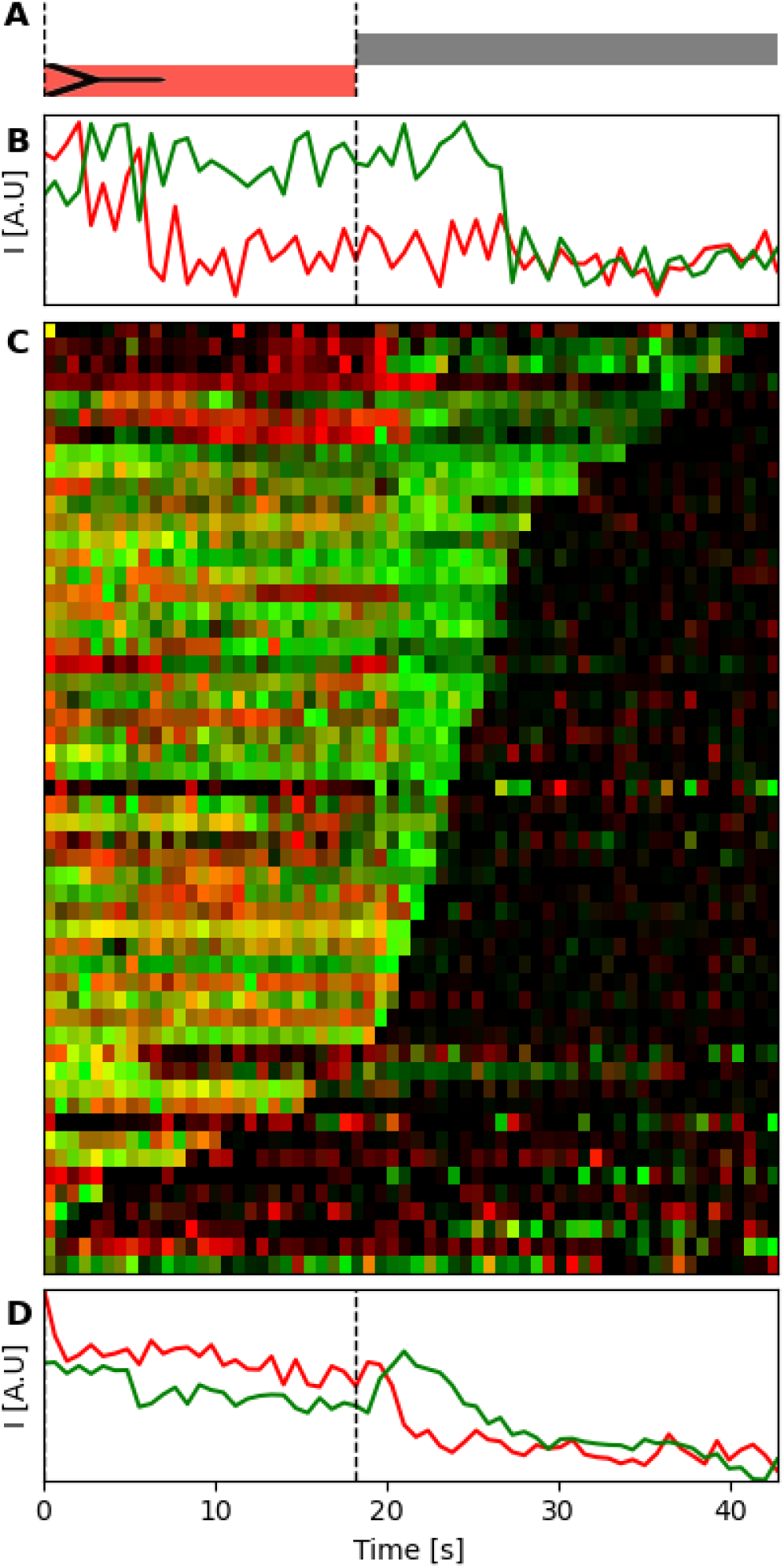
TW rapidly detached from DNA track after removal of ligands (Trp+Co_2+_), as observed by single-molecule FRET. **(A)** Schematic of expected TW position and false-color representation of expected fluorescence signal given solution changes (dashed lines). **(B)** Normalized example single-molecule trace. **(C)** Kymographs of all detected colocalizing single molecules sorted by time of last identified fluorescence. **(D)** Sum of all single-molecule traces in the experiment.

**Extended Data Fig. 10.**
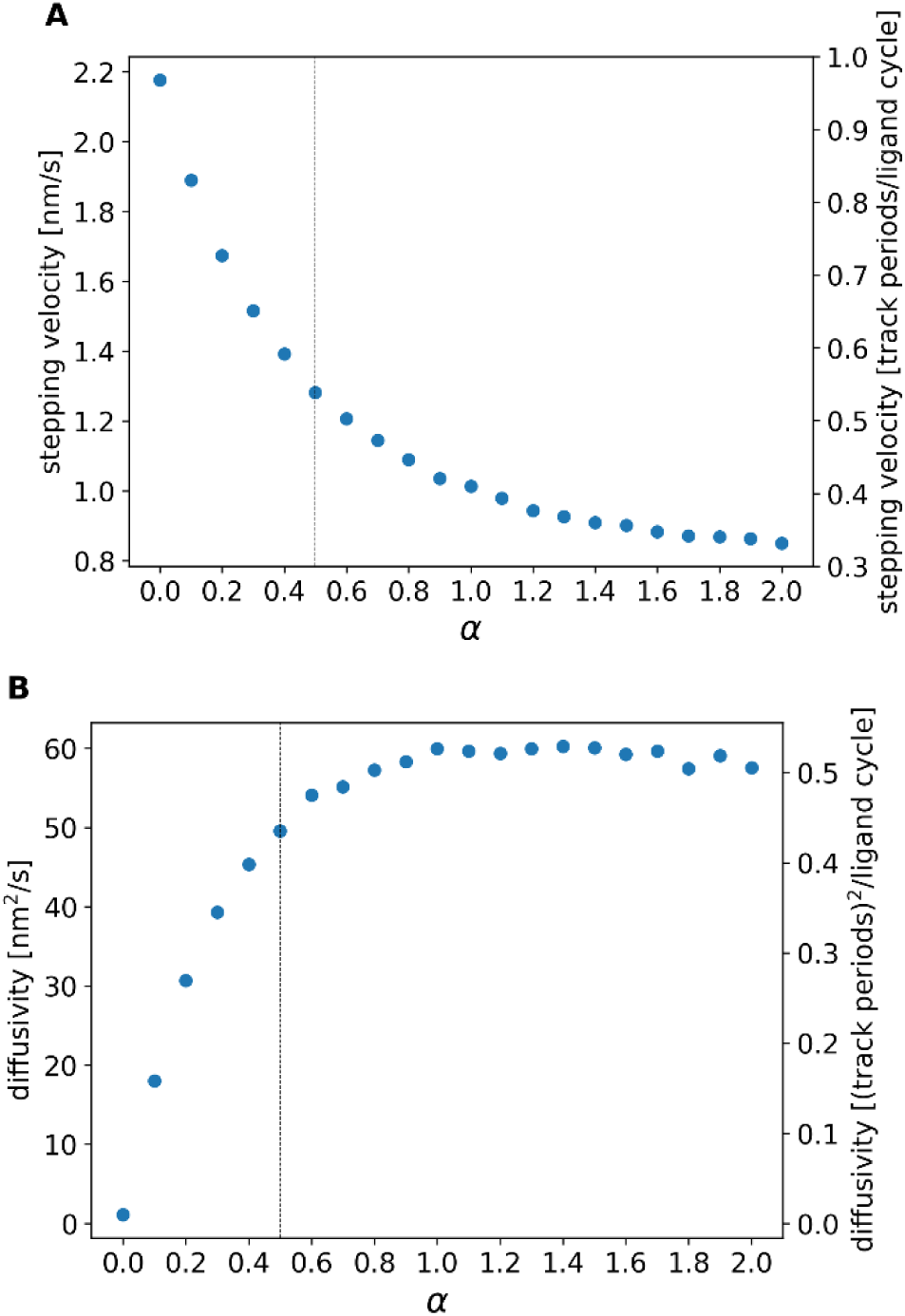
Dynamical characteristics of the stepping motion along the track as a function of the parameter a which controls the relative frequency of overstepping. For a=0 there is no overstepping, a=0.5 corresponds to the situation observed in experiments where 65% of the time TW correctly steps while 35% oversteps (vertical dashed lines), for a=1 correct stepping and overstepping are equally likely, and for a>1 overstepping dominates. **(A)** Average stepping velocity obtained from 500,000 independent realizations of the stepping simulation. For the units of velocity, a track period consists of three binding sites and a ligand cycle is considered a full cycle of three ligand solution changes, so that a velocity of 1 corresponds to ideal stepping. **(B)** Effective diffusion coefficient of the same set of 500,000 trajectories.

**Extended Data Fig. 11.**
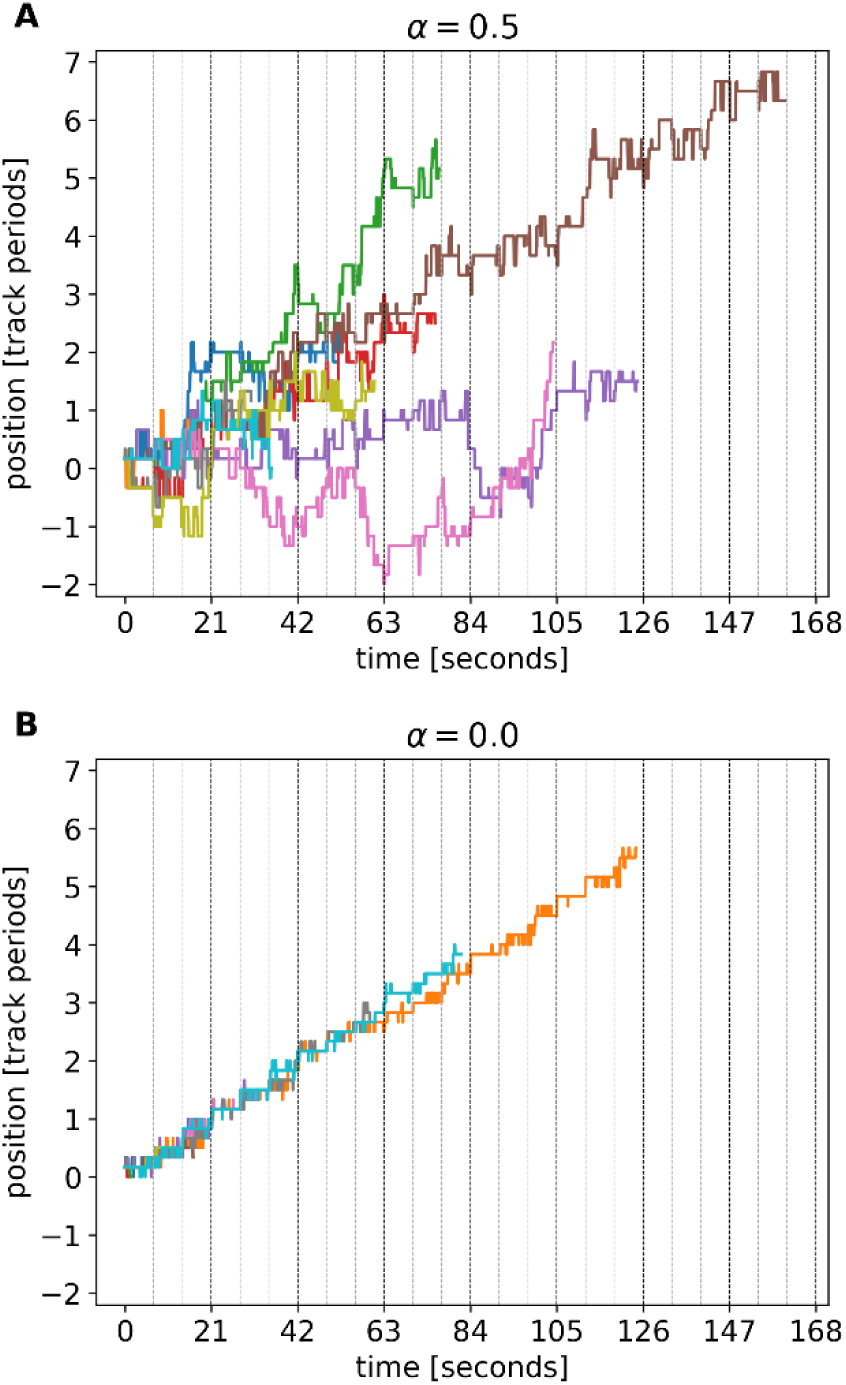
Trajectories of TW stepping along the DNA track from simulations. In each panel 10 independent simulation runs are shown in different colors. The vertical dashed lines indicate the time points at which the ligand combination in the solution is switched. **(A)** Trajectories with a probability of overstepping that reproduces the experimentally observed 65% correct stepping and 35% overstepping. **(B)** Trajectories without overstepping (only correct binding to the track is included in the model). Substeps visible in the traces correspond to binding/unbinding events within the current spatial period on the DNA track.

**Extended Data Fig. 12:**
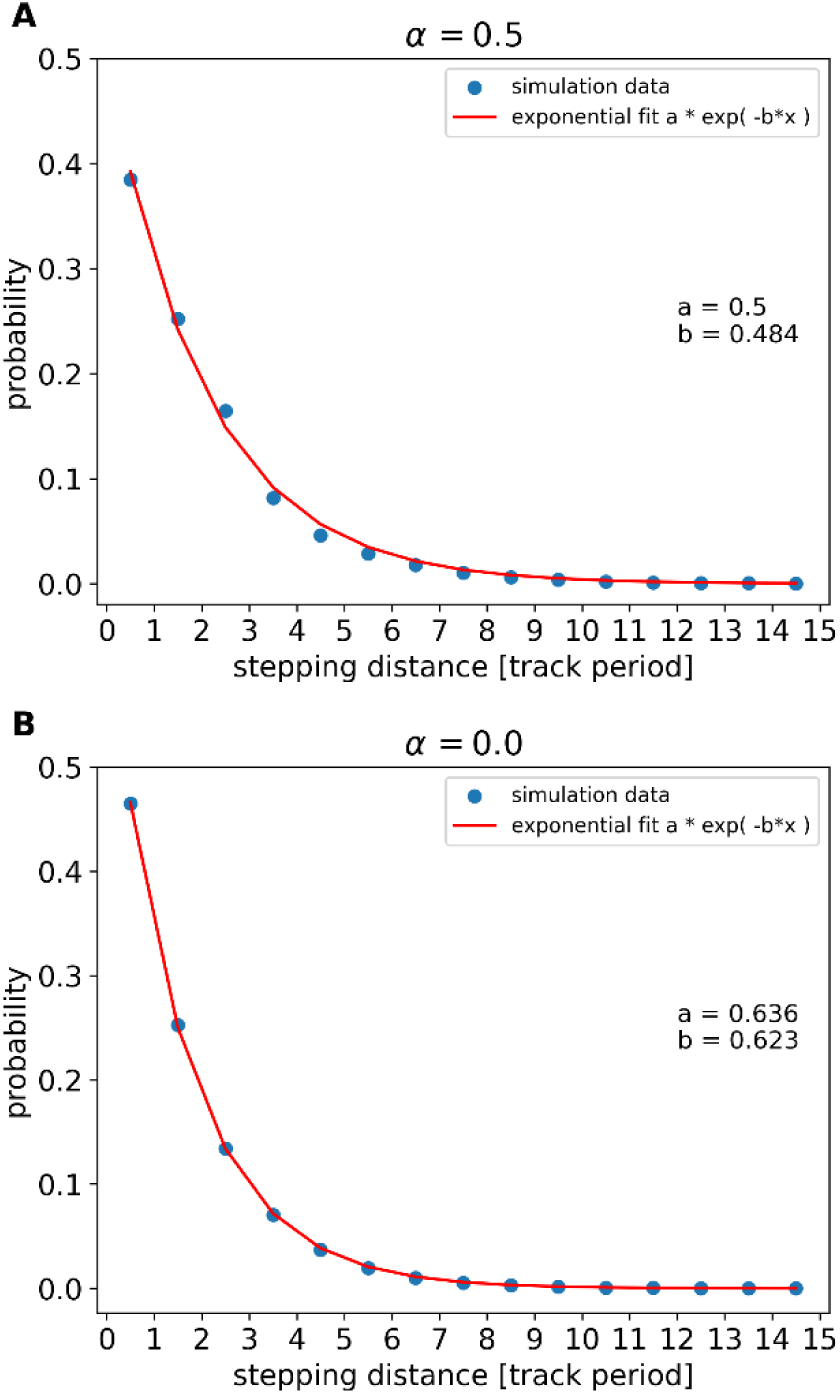
The distances the simulated TW steps along the track before detaching are exponentially distributed. From a fit to the exponential curves, we obtain the average distance covered before detaching from the track (in units of [track periods]). **(A)** (left panel): a=0.5 (experimentally observed frequency of overstepping); 1/e stepping distance is 2.1 track periods ≍ 103 nm. **(B)** (right panel): a=0 (no overstepping); 1/e stepping distance is 1.6 track periods ≍ 78 nm.

**Extended Data Fig. 13.**
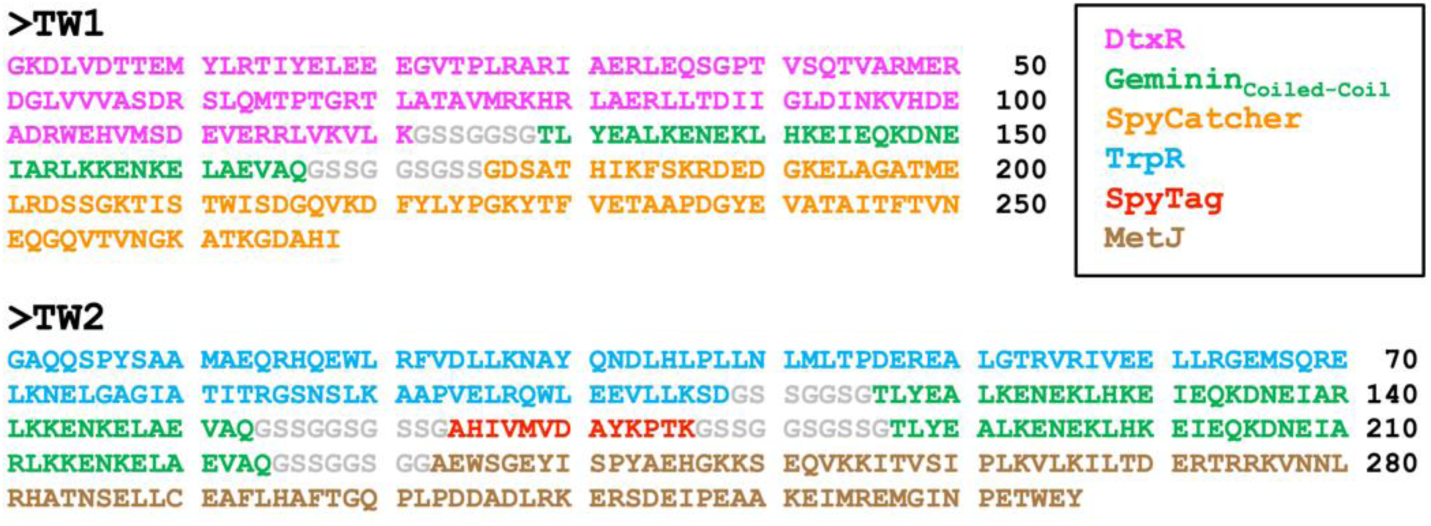
Protein sequences of the TW constructs. Protein domains are colored per the top right inset. Glycine-serine rich linkers are colored grey. TW is formed via the formation of an isopeptide bond between the side chains of K186 in the SpyCatcher domain of TW1 and D170 in the SpyTag part of TW2. The S107C mutation was introduced into the TrpR domain of TW2 to enable labelling of the protein with AF488 via maleimide chemistry.

**Extended Data Fig. 14.**
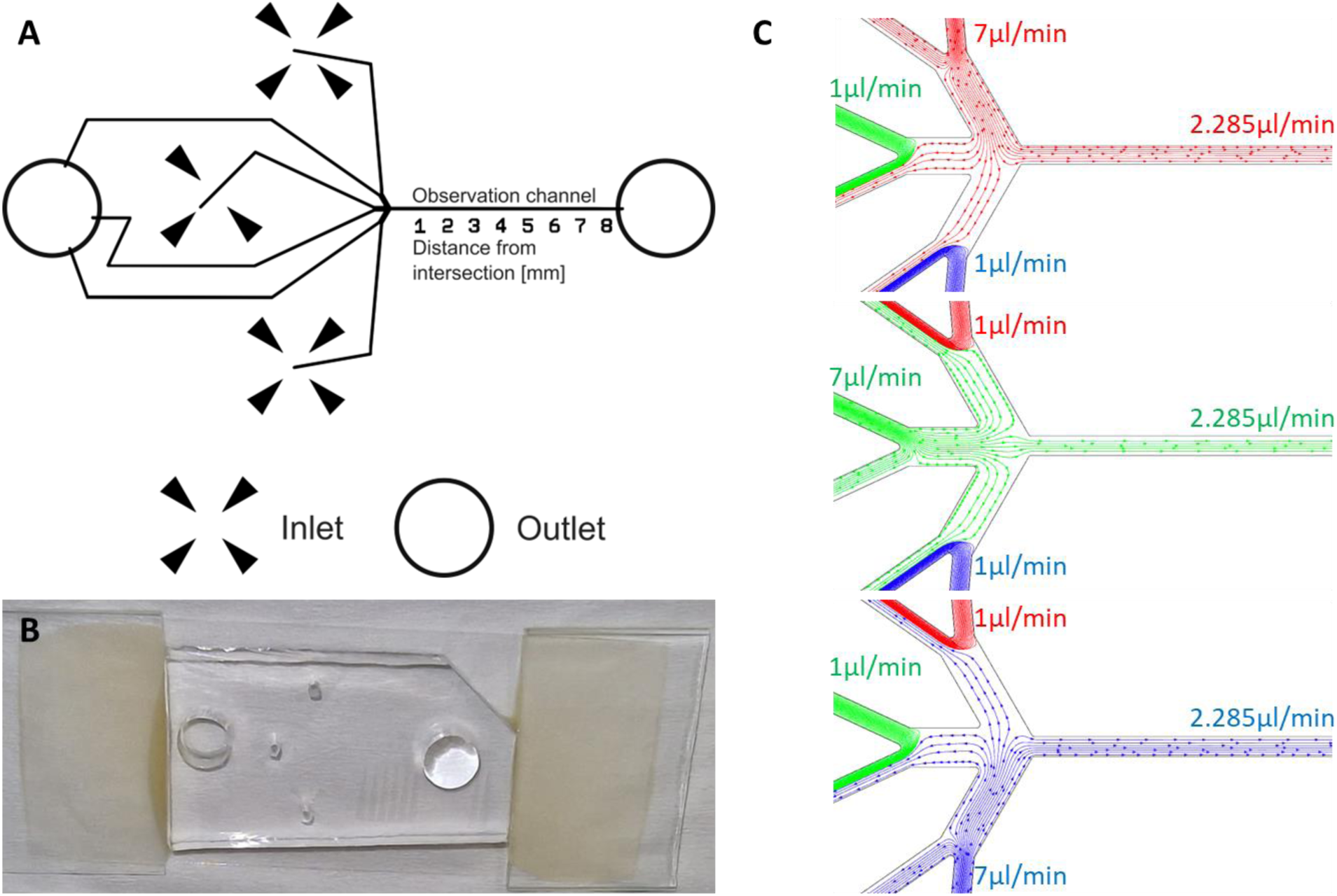
Design of microfluidic device for single molecule TW stepping experiments. **(A)** Locations where the three inlets are punched are marked with arrows pointing inwards. Each inlet is to be connected to a reservoir containing the running buffer with a pair of ligands. Each inlet is connected via an equal length escape channel to a common outlet (circle, far left). Directly following the intersection of the three inlet channels, the main observation channel can be seen labelled with distance in mm from the intersection. The observation channel terminates in another outlet (circle, far right) placed 10 mm from the intersection. **(B)** Photo of assembled microfluidic device bonded to a coverslip including glued glass slide fragments for support and mounting. **(C)** 2D finite element simulation (incompressible, laminar flow) of flow rates in the microfluidic device for three fluids (red, green and blue) injected into the device inlets, one at 7 µL/min (“on” channel) the other two at 1 µL/min (“off” channel). The fluid injected at 7 µL/min (Top: red, Middle: green, Bottom: blue) propagates in the observation channel at 2.285 µL/min while the other two fluids are directed to their respective “escape” channel.

**Extended Data Fig. 15.**
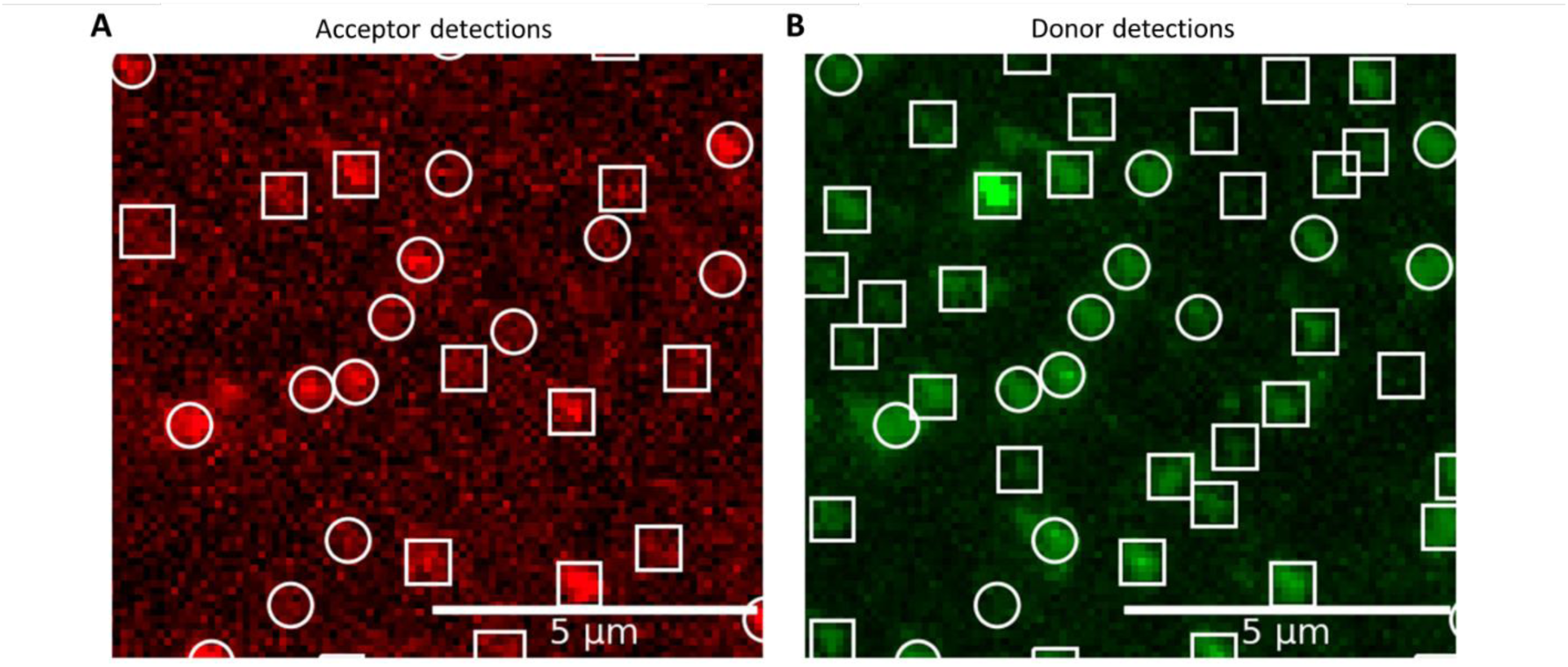
Example of detected bright spots. A small field of view from a normalized maximum intensity projection of the data from **(A)** the FRET acceptor channel and **(B)** the donor channel from Experiment 1. All bright spots detected by the algorithm are outlined with white squares or circles. Squares denote those detected spots that were identified as not colocalizing in Donor and FRET channels. Circles denote those detected spots that were identified as colocalizing in Donor and FRET channels.

**Extended Data Fig. 16.**
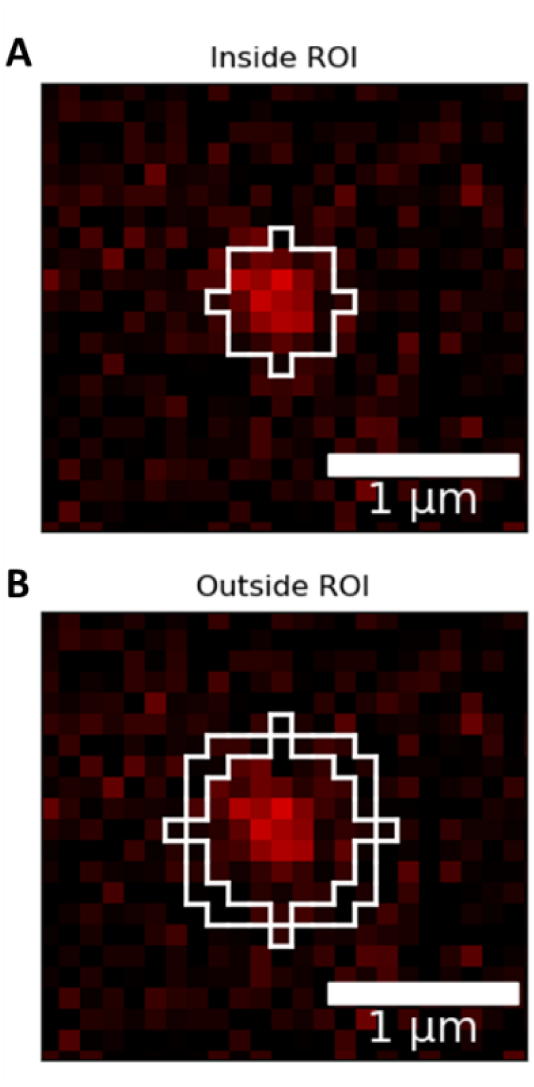
Regions of interest created for single-molecule analysis. **(A)** shows the pixels used to determine intensity of a spot, and **(B)** shows the surrounding pixels whose intensity is used for background removal. The final background subtracted intensity of the molecular fluorescence signal in each frame is calculated by averaging the intensity of all pixels in the region of interest in **(A)** and subtracting the median in **(B)**.

**Extended Data Fig. 17.**
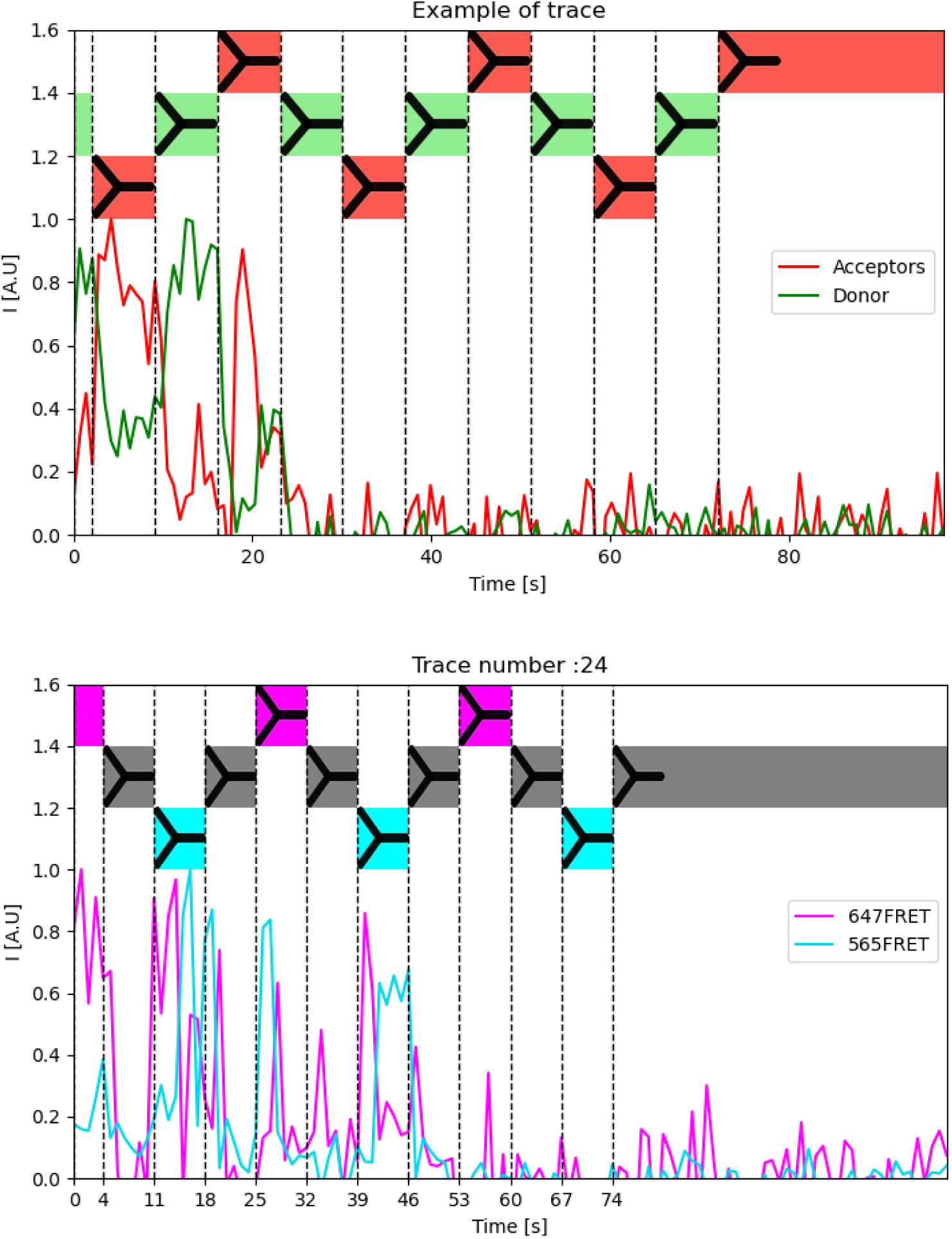
The resulting traces after background subtraction and normalization. Top: Experiment 1. Red is the FRET acceptor signal, green the donor. Black dashed lines indicate times of buffer changes. Bottom: Experiment 2. Magenta is the FRET acceptor signal from the red acceptor (top-*trpR* site) and cyan is the FRET acceptor signal from the yellow acceptor (bottom-*trpR* site). Black dashed lines indicate buffer switches.

**Extended Data Fig. 18.**
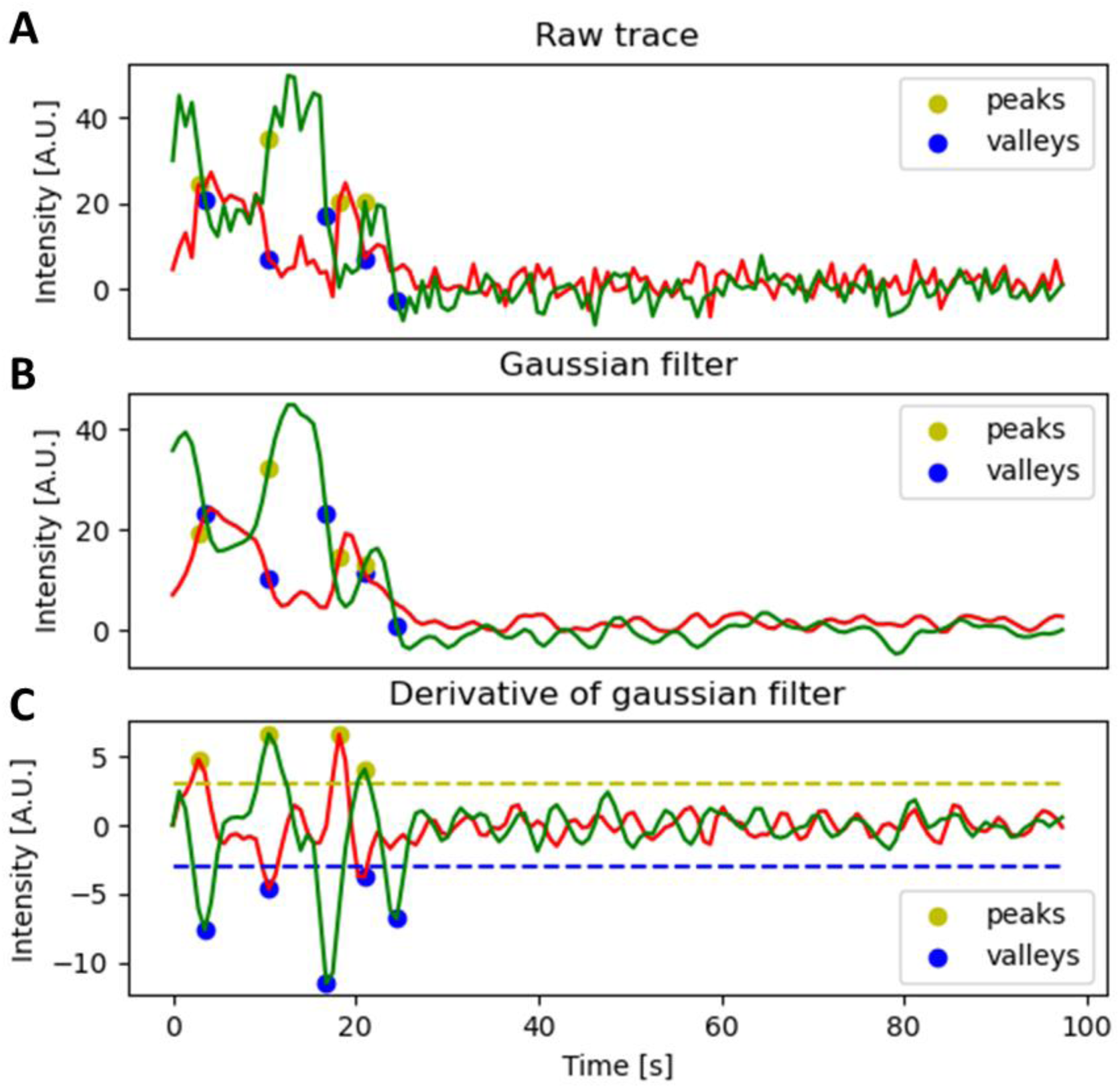
Outline of the edge detection algorithm. **(A)** shows the raw traces, **(B)** the raw traces after Gaussian filtering with standard deviation 1.2 and **(C)** shows the derivative of **(B)**. The threshold for peak detection is indicated by a yellow dashed line and the threshold for valley detection is indicated by a blue dashed line. In all panels, the detected rising edges are marked with a yellow dot and detected falling edges with a blue dot.

**Extended Data Fig. 19.**
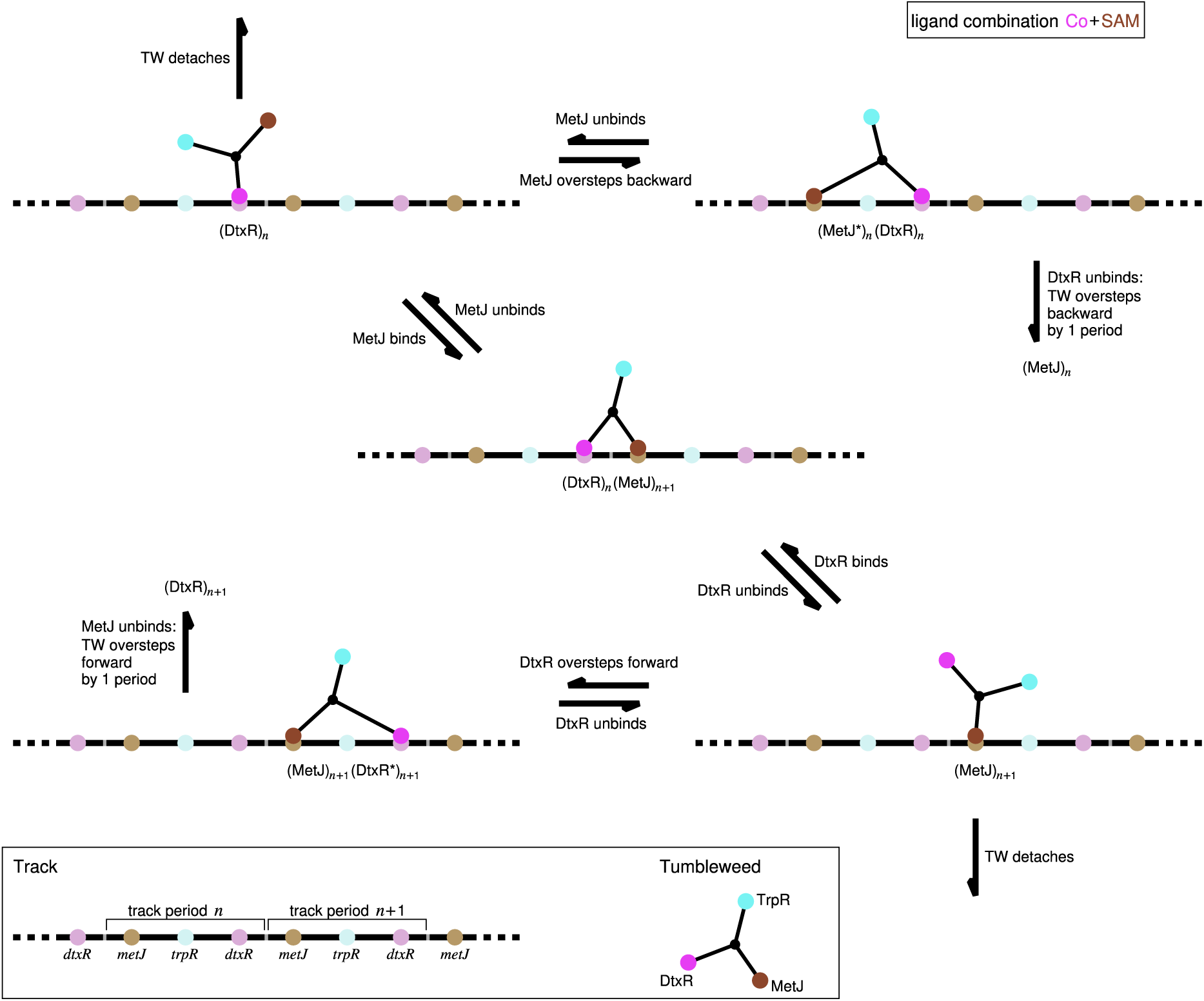
Illustration of states and transitions modelled by the rate equations. Lower panel: The DNA track, extending ad infinitum, is sketched by a horizontal black line with colored dots, which label the different cognate binding sites on the track (compare Fig. 1B in the main text); two track periods are marked as *n* and *n*+1. The TW molecule is depicted in a star-like manner with three colored dots representing the repressor proteins TrpR (cyan), DtxR (magenta) and MetJ (brown), see Fig. 1 in the main text. Main panel: The five different possible states in the presence of the ligand combination Co^2+^ + SAM, including intended binding events and unintended overstepping events. The states are labelled by the repressor(s) bound to the track and the track period they are bound to, e.g., (DtxR)*_n_*(MetJ)*_n_*_+1_ indicates that DtxR is bound to track period *n* and MetJ to track period *n*+1 (this is the state in the middle); an asterisk at the repressor label indicates an overstepped configuration. The arrows indicate transitions between these different states. For the chosen ligand combination, there are no states in which TrpR is bound, or transitions towards such states, because we assume that TrpR can only bind in the presence of its ligand Trp. The sequence of events from the state (DtxR)_n_ (upper left corner) to the state (MetJ)_n+1_ (lower right corner) via the state (DtxR)*_n_*(MetJ)*_n_*_+1_ in the middle corresponds to the desired forward stepping behavior from one track period into the next. The transitions from the states (MetJ*)*_n_*(DtxR)*_n_* and (MetJ)*_n_*_+1_(DtxR*)*_n_*_+1_ marked “TW oversteps” end up in the states (MetJ)_n_ and (DtxR)*_n_*_+1_ (not shown), which are equivalent to (MetJ)_n+1_ and (DtxR)*_n_* but shifted by one track period backward and forward, respectively.

**Extended Data Fig. 20.**
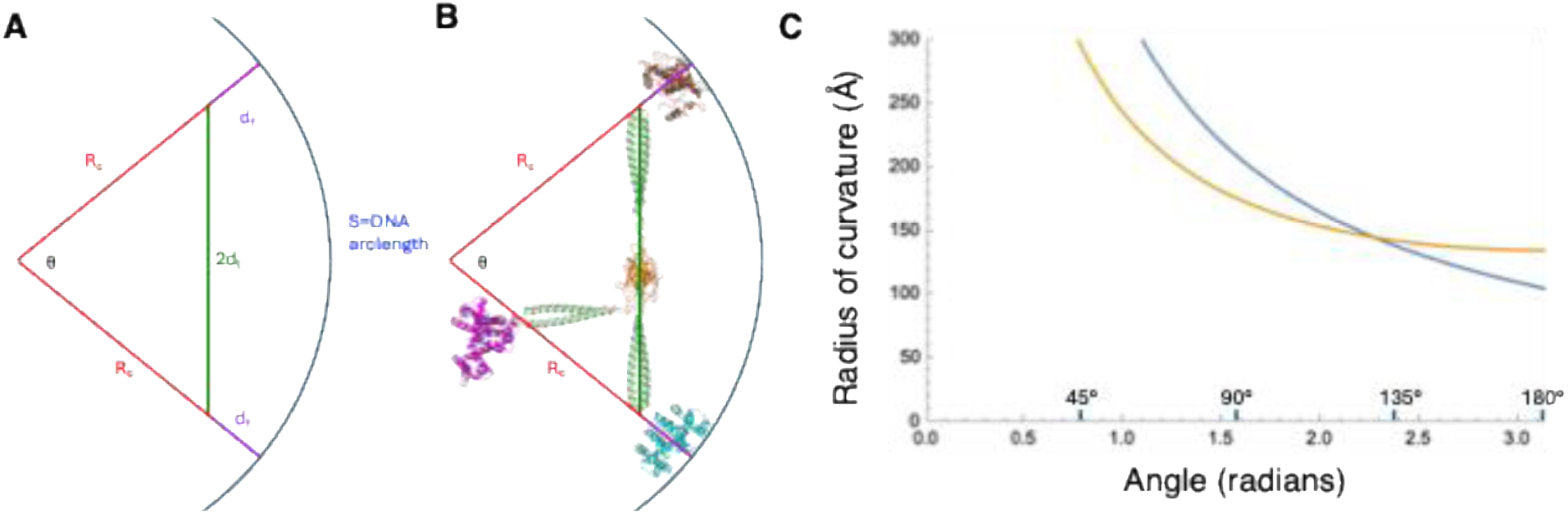
Model for TW overstepping due to DNA flexibility. **(A)** The geometry of DNA bending into a uniform circular arc. TW is represented by two stretched legs, total length *2d_l_* that connect to DNA via two repressor feet, each of length, *d_f_*, that bind to DNA in a perpendicular fashion, each tracing a radius of curvature. **(B)** An overlay of a model of fully stretched TW on the geometric figure from (**A**). (**C**) Plot of the equation for the radius of curvature describing the DNA geometry (blue) and the TW geometry (orange) as a function of angle. The intersection of these two curves marks a simultaneous solution to the two equations.

## Supplementary

## Supplementary Discussion A

### Single molecule FRET: Control Experiments

The effect of the fluid exchange in the absence of ligand changes was tested by performing the same biochemical preparation, imaging and mechanical flow switching as in the stepping experiments, but instead with Co^2+^+SAM buffer in each of the three microfluidic reservoirs. Extended Data Fig. 7 shows traces obtained using a single-molecule FRET analysis. In this experiment the TW is never expected to bind to a site labelled with a FRET acceptor, so the analyses for colocalization was relaxed by removing the requirement for colocalization with FRET intensity. Instead, relying only on colocalization with detections in the acceptor excitation channel. Keeping the FRET requirement removes all molecules from the analysis as expected. Otherwise, analysis was performed as in the above Methods section. Solutions were changed between identical Co^2+^+SAM buffer 3 times and then changed to a buffer containing no ligands at all as indicated by Extended Data Fig. 7A. Extended Data Fig. 7B shows an example of these traces where TW dissociates from the DNA after the removal of ligands. All traces are shown as overlayed kymographs in Extended Data Fig. 7C where green is TW donor signal and red is *trpR* site FRET. The kymograph shows no FRET nor any reaction to the changes in flow at all. At the point at which all ligands are removed (Extended Data Fig. 7A) a slower than anticipated rate of signal loss is observed (Extended Data Fig. 7B). This is due to the relaxed colocalization criteria allowing non-specifically bound nearby TW and DNA and potentially non-functional TW:DNA complexes to be included in the analysis. This is particular to this control experiment where we do not have a FRET signal as an analysis criterion.

Similar detachment experiments were performed with Trp+Co^2+^ (Extended Data Fig. 8) and SAM+Trp, (Extended Data Fig. 9), with a single solution change to the ligand-free solution. When changing to a ligand-free solution, the rate of TW detachment increases sharply for both of these ligand-pair conditions.

## Supplementary Discussion B

### Simulation of TW Stepping

The single-molecule FRET data in Fig. 4 clearly demonstrate that TW can step bi-directionally on a short track consisting of four binding sites, but does not allow us to directly and experimentally evaluate performance parameters such as processivity and average speed on a long track. However, experimentally measured binding and stepping characteristics allow us to predict this performance using a model.

To analyze potential stepping behavior of TW along an extended track we devised a coarse-grained model based on Master equations governing the binding and unbinding kinetics of the three TW feet and the track. The track is represented by the periodic, unlimited sequence of binding sites …*-dtxR-metJ-trpR-dtxR-metJ-trpR-dtxR-metJ*-… (Extended Data Fig. 19), mimicking the experimental setup used in Fig. 4. The kinetics of the three individual feet are described by three independent rate equations. The rate equation for each foot describes the transitions between an unbound state and bound states, in which the foot is bound specifically to its matching binding site on the track, e.g., the DtxR foot can only bind to *dtxR* etc., while non-specific binding is neglected, supported by the SPR results in Fig. 3A. Since these equations govern transitions between states (not concentrations), the corresponding rates have units [1/time] and are the inverse of the average waiting time between transitions. The values of these rates depend on the ligands in solution (see Extended Data Table 8), which are cycled periodically through the combinations Trp + Co^2+^, Co^2+^ + SAM, SAM + Trp, with a period of 21 seconds (i.e. the solution is switched every 7 seconds). Since there are always two ligands present at any given time, the rate equations for the individual feet become effectively coupled.

As an example, the set of possible states and the considered transitions between them are shown in Extended Data Fig. 19 for the ligand combination Co^2+^ + SAM. In particular, Extended Data Fig. 19 illustrates that we set up the rate model for each foot to distinguish between two distinct bound states, representing binding of a foot to its intended site (adjacent to the other bound foot) and unintended binding to the non-adjacent binding site (overstepping). For the other ligand combinations, Trp + Co^2+^ and SAM + Trp, analogous state combinations and transitions are realized by the rate equations.

The dynamics of each foot is thus driven by four rates for the transitions between its unbound state and two distinct bound states, which all depend on the foot-specific ligand being present in the solution or not. Fixing the explicit values for these rates in the model is guided by the experimental findings and is described below.

The rate processes for the three TW feet, their dependence on the ligand combination present in solution, and the 21-second cycle of the solution through three ligand combinations were implemented in a custom-built C++ code. A simulation run started with two feet bound on the track in the presence of the corresponding ligand combination, and ran until the TW detached from the track, *i.e.*, until none of the feet was in a bound state. The sequence of binding and unbinding events recorded during such a run for all three feet was translated into the corresponding stepping motion of the TW along the track. This resulted in a trajectory of the TW showing the position of its center (in units of track periods) as a function of time (in seconds). Examples of such trajectories are shown in Extended Data Fig. 11.

Repeating simulations for individual trajectories many times gives access to the distribution of observables with which the expected stepping performance of the TW along an extended track can be characterized. We focus on the stepping distance as a function of time (giving the stepping velocity, Extended Data Fig. 10A), the variance of the trajectories as a function of time (giving the effective diffusion coefficient, Extended Data Fig. 10B), and the distance covered before the TW detaches from the track (Extended Data Fig. 12).

The specific values for the various rates of the individual TW feet in our model were chosen as follows. We first focus on the rates describing (un-)binding between the TW repressor feet and the intended binding site (adjacent to the other bound foot): The off-rates for each repressor in the presence and absence of its ligand were adapted from the SPR experiments, see Extended Data Table 8 (compare to Extended Data Table 3), *i.e.*, the off-rates are characteristic for the type of repressor (TrpR, DtxR, MetJ). To choose the on-rates, we assumed that (re-)binding to the track is predominantly determined by the geometrical arrangement of the track and the binding sites, and thus can be assumed to be independent of the repressor type. For this reason, we took on-rates to be identical for all feet. Moreover, neglecting non-specific binding, the on-rates are all zero in the absence of the foot’s ligand (see Extended Data Table 8). In the presence of ligands, we adjusted the value of the on-rates such that the three *double-ligand* off-rates measured in the SPR dissociation experiments for the ligand combinations Trp + Co^2+^, SAM + Co^2+^, and SAM + Trp (Extended Data Table 3) were best reproduced. To do this, we ran 10^6^ simulations for each pair of feet and corresponding ligands, with the assumed value of the on-rates, until neither of the feet was bound . From the exponential distribution of the so obtained “survival times” we extract the effective dissociation rate; the results are listed in Extended Data Table 8 (compare to Extended Data Table 3 for the experimental values).

For unintended binding to the non-adjacent site (overstepping), we assumed the off-rates to be identical to the ones for unbinding from the intended binding site, because the off-rates are characteristic for the type of repressor, no matter where on the track it is bound. In the spirit of our assumption that the binding process is predominantly geometry-determined, the on-rates are again all the same for the different feet, but in general different from the on-rates to the intended binding site (different geometry). We encoded the difference in a factor a that multiplies the on-rates for intended binding:

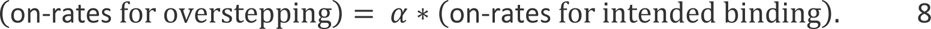

For a=0, there is no overstepping, 0<a<1 implies that overstepping is less likely than intended binding, while a>1 indicates that overstepping is more likely; for a=1 intended binding and overstepping are equally likely. The relative weight of overstepping is calculated by comparing its rate to the total rate of binding, a/(1+a). Hence, for a=0.5 we find 1/3=33%, reproducing the experimentally observed frequency of overstepping of about 35%.

## Supplementary Discussion C

### Can DNA bend sufficiently to facilitate overstepping?

The inherent flexibility of the DNA track may be responsible for the observed overstepping of TW. To test this possibility, we used a simple physical model to determine the energetic cost of bending the DNA sufficiently to allow overstepping, *i.e.*, for two repressor feet to bind sites separated by an intervening empty site.

If DNA is uniformly flexible, then it can follow a circular arc described by a radius of curvature, *R_c_*, an arc length, *s*, and the angle subtended by the arc, *θ*, where

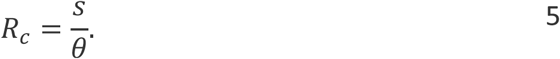

The largest possible reach of two TW feet occurs when the two legs (each of length *d_l_*) are straight and each repressor foot (additional length *d_f_*) binds to DNA perpendicularly (thus aligning the foot’s axis with the local radius of curvature) (Extended Data Fig. 20A,B; similar to Model 2 in Fig. 2C). For an angle *θ* subtended by the feet, this configuration of TW can be described by:

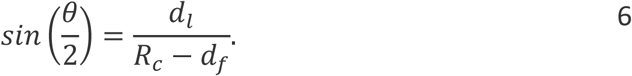

From our model of TW, we approximate: 𝑑_𝑙_ = 100Å and 𝑑_𝑓_ = 35Å. From our model of the DNA track, the arc length between two non-consecutive repressor binding sites is approximately 𝑠 = 330Å. Using these parameters, we can plot equations 5 and 6 for radius of curvature, *R_c_*, as a function of the angle *θ* subtended – the equation 5 describing the DNA arc and equation 6 describing TW. The intersection of these two curves represents the solution of these two geometric equations (Extended Data Fig. 20C), where 𝑅_𝑐_ ≅ 145Å and 𝜃 ≅ 130°.

By modelling DNA as a rigid rod of length *s*^61^, we can estimate the energy, *ΔU*, required to bend DNA sufficiently to satisfy this geometric solution:

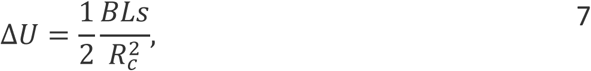

where *B* is the bending stiffness, 𝐵 = 50𝑛𝑚. 𝑘_𝐵_𝑇 ^61^. This gives Δ𝑈 = 3.9𝑘_𝐵_𝑇, where *k_B_* is the Boltzmann constant. Thus, it would not require a large thermal fluctuation to achieve the required bending of the DNA that would facilitate overstepping (along with flexibility in the TW linker regions). This bending could be reduced by supporting the DNA track in a straight configuration.

## Notes

### Competing Interest Statement

The authors have declared no competing interest.

### Summary of Updates

Edited to align with style of new journal, clarified abstract, introduction and discussion sections.

